# Ancestral Body Plan and Adaptive Radiation of Sauropterygian Marine Reptiles

**DOI:** 10.1101/2022.04.25.489368

**Authors:** Wei Wang, Qinghua Shang, Long Cheng, Xiao-Chun Wu, Chun Li

## Abstract

Mesozoic was the age of reptiles that not only occupied the land and sky but also adapted to the sea. Sauropterygia, from turtle-like placodonts, lizard-like pachypleurosaurs, and predatory nothosaurs to long-necked pistosaurs including plesiosaurs, is one of the most predominant lineages among secondarily aquatic reptiles. However, we know little about the early evolution of sauropterygians involving the morphological gaps among aforementioned sub-groups and their ancestral body-plan, because the Early Triassic fossil remains are scarce shortly after their origin. Here we report a skeleton from the Early Triassic in South China, representing the oldest known complete specimen related to Sauropterygia. It can be referred to *Hanosaurus hupehensis* and shows a mosaic morphology combining the characters of multiple sauropterygian sub-lineages. We constructed an updated character matrix obtaining the hitherto most comprehensive phylogeny of Triassic sauropterygians, and *Hanosaurus hupehensis* is stably resolved as the basal-most member of the Sauropterygiformes (clade nov.). Sauropterygians were previously considered using limb propulsion for the movement that is distinguishable from ichthyosaurs and other marine reptiles, but this skeleton reflects an ancestral body plan of sauropterygians unexpectedly developing an elongate trunk and four short limbs, being different from most sauropterygians but more convergently similar to the basal members of other marine reptilian lineages in body shape. After this convergence, we confirm the rapid adaptive radiation of sauropterygiform reptiles following the end-Permian mass extinction. Our results provide an evolutionary framework of sauropterygian marine reptiles and highlight the integrations of both convergences and divergences when an emerging animal lineage arises.

## RESULTS

Marine reptile is a great model for evolutionary study by exhibiting secondarily aquatic adaptations^1^. Sauropterygia^2^ is one of the most diversified and dominant marine reptilian groups having been studied for about two hundred years. Based on abundant complete skeletons of sauropterygians and other significant marine reptiles^3,4^ discovered during recent two decades, especially from southern China^5^, we could have better glimpses of sauropterygian origin and early evolution. Here we report a nearly complete skeleton referred to *Hanosaurus hupehensis*^6^. In this research, we aim to resolve the relationship within sauropterygians with related marine reptiles such as saurosphargids, illuminate the very early history, and evaluate the adaptive radiation of sauropterygian and allied reptiles.

### Systematic paleontology

> Diapsida Osborn, 1903
>
> Sauropterygiformes clade nov.
>
> *Hanosaurus hupehensis* Young, 1972

#### Holotype

Institute of Vertebrate Paleontology and Paleoanthropology (IVPP) V 3231, a skull (Supplementary Fig. 1) in dorsal view and a partial postcranial skeleton (Supplementary Fig. 2) in ventral view^6,7^.

#### Type Locality and horizon

Songshugou, Xunjian Village, Nanzhang County, Hubei Province, China; dolomitic limestone (Member II) of the Jialingjiang Formation, Olenekian, Early Triassic^5^.

#### Referred specimens

IVPP V 15911, a nearly complete skeleton in ventral view (Fig. 1, Supplementary Fig. 1 and 2).

**Figure 1.**
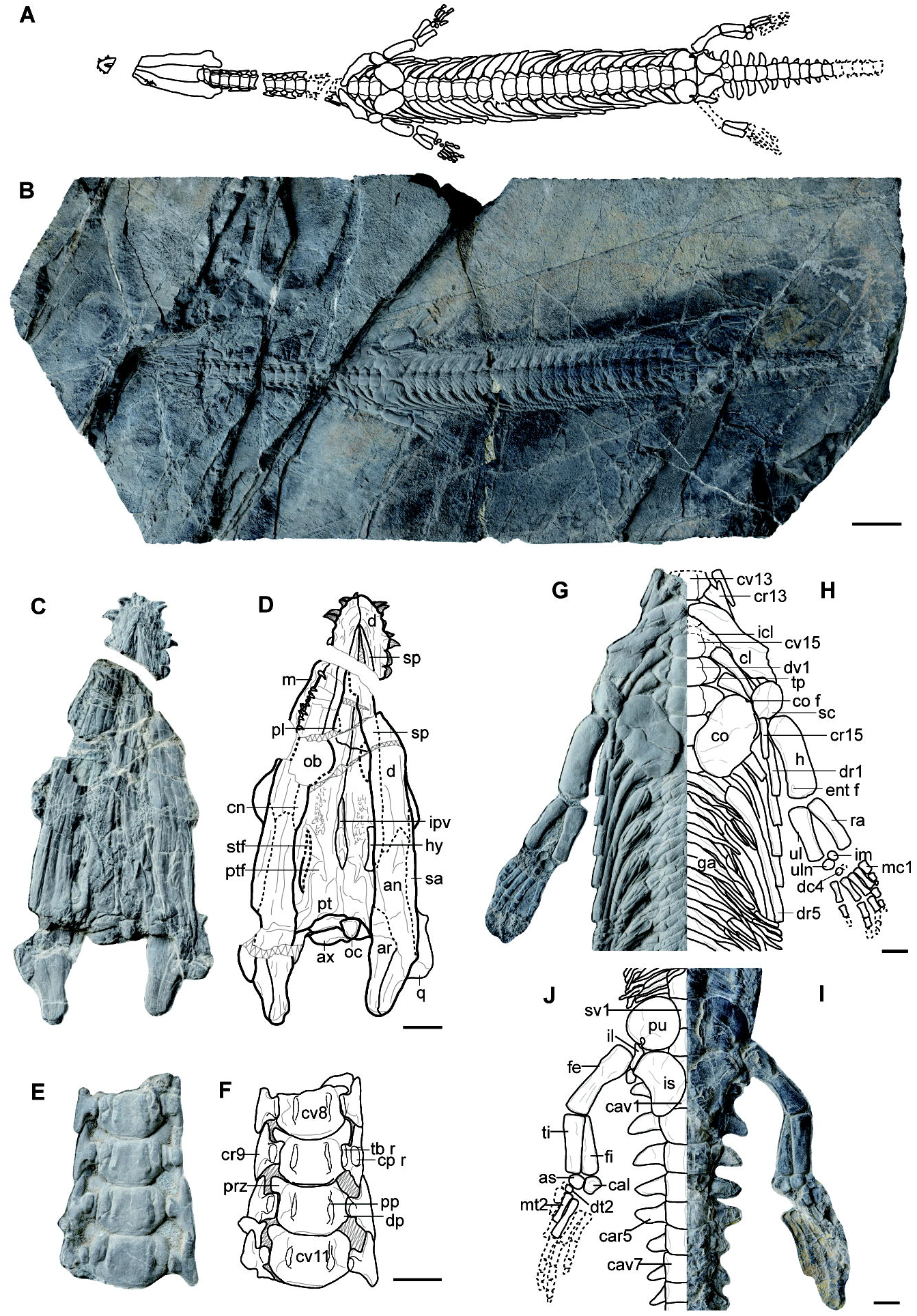
The first complete skeleton of *Hanosaurus hupehensis*. IVPP V 15911. (a-b), new specimen preserved in ventral view; (c-d), skull; (e-f), the medium region of cervical vertebrae; (g-h), the right side of the anterior region of the trunk with the pectoral girdle and the right forelimb, the line drawing is a mirror of the opposite side along the midline; (i-j), the left side of the sacral region and the anterior region of the tail, with the pelvic girdle and the left hind limb, the line drawing is a mirror of the opposite side along the midline. (scale bar=5 cm in a-b, =1 cm in all the others; see abbreviations in Supplementary Information)

#### Locality and horizon

Yingzishan, Yuanan County, Hubei Province, China; dolomitic limestone (Member II) of the Jialingjiang Formation, Olenekian, Early Triassic.

#### Amended Diagnosis

A medium-sized sauropterygiform reptile with the exclusive combination of snout without constriction; supratemporal fenestrae smaller than orbits; nasals longer than frontals; mandibular articulation far posterior to the level of occiput, with retroarticular processes well-developed; anterior teeth short and conical; posterior teeth constricted at the base, with the lingual surface of tooth crown slightly concave; cervical ribs with evident anterior processes; dorsal ribs pachyostotic; coracoid not waisted; pubis extremely round; ischium kidney-shaped; no distinct thyroid fenestra between pubis and ischium; and minimally four carpals and three tarsals.

### Description

This new specimen, IVPP V 15911, is entirely articulated and exposed in ventral view, with a total length of 79.4 cm (Fig. 1 A, B) being comparable in size to IVPP V 3231, the holotype of *Hanosaurus hupehensis*. The cranial shape is of an elongate triangle (Fig. 1 C, D). The snout is neither pointed nor blunt without lateral constriction. A pair of large pterygoids meet along the midline suture anteriorly and posteriorly leaving a narrow interpterygoid vacuity. In the holotype, the preorbital and postorbital regions of the skull are nearly equal in length, and the supratemporal fenestra is smaller than the orbit (Supplementary Fig. 1), which is similar to saurosphargids^8^, pachypleurosaur^2^, and pachypleurosaur-like reptiles^9^ but on the contrary to derived eosauropterygians such as nothosaurs and pistosaurs^2^. The same positions and proportions of orbits and supratemporal fenestrae can be tentatively identified in our new specimen on the right side of the skull from the ventral view. The short mandibular symphysis is mainly contributed by the dentary. The retroarticular process is well developed as in the holotype, with a blunt termination.

There are at least 6 dentary teeth exposed on the anterior portion of the right dentary and more than 10 maxillary teeth visible on the right side (Fig. 1 C, D). There is neither obvious diastema between premaxillary and maxillary teeth nor prominently enlarged fangs. All the teeth are short, sharp, and likely thecodont. The anterior teeth are generally coniform, slightly procumbent; they are relatively larger than the posterior teeth and possess longitudinal striations. The lateral teeth are smaller, mildly overbite, and show relatively smooth surfaces of the crowns with slight constrictions at the bases of the crowns (Fig. 1 C, D; Supplementary Fig. 2). Additionally, same as those in the type specimen (Supplementary Fig. 1 and 2), the crowns of the lateral teeth are slightly lingually concave showing leaf-shaped outlines, which resemble those in saurosphargids^8,10^.

There are 15 cervical, 25 dorsal, and 3 sacral vertebrae, and at least 16 caudal vertebrae in IVPP V 15911. The number of cervical vertebrae here in *Hanosaurus* (15 in total with 13 anterior to the pectoral girdle) is fewer than all Triassic eosauropterygians. We identify the last cervical vertebra according to the corresponding rib not articulated with the sternum^11^ instead of the position anterior to the pectoral girdle. Therefore, the previously reported cervical counts^12^ should be revised as about 20 in *Neusticosaurus* and 18 in *Serpianosaurus* based on our first-hand observation, which develops fewer cervical vertebrae than other Triassic eosauropterygians but more than *Hanosaurus*. Including the ribs, the entire vertebral column is conspicuously pachyostotic. The centrum is wider than long and anteriorly narrow and posteriorly wide in ventral view (Fig. 1 A, B, G, H). The cervical rib possesses a distinct anterior process in vertebrae 2 to 13 (Fig. 1 E-H; Supplementary Fig. 2 I-L). The last cervical rib is elongate, single-headed, and nearly as long as the first dorsal rib, but its shaft is thinner than the latter (Fig. 1 G, H). The anterior caudal ribs are anteroposteriorly broader than the gap width between the ribs and turn sharp laterally as in the holotype (Supplementary Fig. 2 M-P). The caudal ribs gradually become short posteriorly and disappear at the 12th caudal vertebra. The lateral elements of the gastralia are straight, and the median elements are angulated, each with a single lateral process (Fig. 1 G, H).

The clavicle is relatively broader than that of most pachypleurosaur-like sauropterygians^2,13^ and nothosaurids^14^, and its expanded posterodorsal portion does not have an anterolateral process (Fig. 1 G, H). The clavicle anteriorly contacts the scapula (Fig. 1 G, H; Supplementary Fig. 2 A, B). The scapula develops a distinct ridge along the midline of the ventral surface and a pit at the posterior end of this ridge. The coracoid is oval with a coracoid foramen open at the suture with the scapular, resembling that in saurosphargids^8,10^, placodonts^2^, and even hupehsuchians^15^ and thalattosaurs, but distinctly different from that in any eosauropterygian taxon. The ilium is not entirely exposed but it is evidently small, much smaller than other girdle elements (Fig. 1 I, J). In both the new specimen and the holotype, the pubis is plate-like and extremely round (Fig. 1 I, J; Supplementary Fig. 2 M-P) being reminiscent of that in saurosphargids and placodonts but different from pachypleurosaur-like sauropterygians and nothosaurids. An open, drop-shaped obturator foramen is present in the pubis near its suture with the ischium as in the holotype (Fig. 1 I, J; Supplementary Fig. 2 M-P). The ischium is kidney-shaped, with an incurved lateral edge and a convex medial margin.

The humerus is nearly straight, with a slightly curved posteromedial margin (Fig. 1 G, H). It is almost uniform, column-like in outline. The ectepicondylar groove is absent, and an entepicondylar foramen is present. The radius and the ulna are equal in length and similar in width. The proximal end of the ulna is slightly expanded. There are at least four roundish carpal ossifications including two large elements, the ulnare, and the intermedium. The femur is also straight and its posteromedial margin is concave; it is as long as but its shaft is thinner than that of the humerus (Fig. 1 I, J). The internal trochanter of the femur is weakly developed but evident, and the intertrochanteric fossa is distinct but reduced. Both the tibia and fibula are straight and nearly equal in length, whereas the former is slightly wider than the latter. At least three tarsals are present, which are identified as the astragalus, the calcaneum, and a distal tarsal abutting to the metatarsal I. Both the manus and the pes are small relative to the body size and do not show a fin-like or a paddle-shaped modification.

### Phylogeny

To investigate the relationship among early sauropterygians, we constructed a new morphological character matrix, which is largely improved compared to previous studies^16-18^ (see Materials and Methods, and Supplementary Information), involving 62 (more than 40 sauropterygian taxa) and 181 characters mostly observed by us on specific specimens. To further examine whether our new specimen can be referred to *Hanosaurus hupehensis*, we coded the holotype specimen of *Hanosaurus hupehensis* (IVPP V 3231) and the new specimen (IVPP V 15911) respectively. Results from both the original matrix of Neenan et al.^16,19^ and our new matrix indicate that these two specimens stably form a single clade (Supplementary Fig. 3, 5). Considering their diagnostic morphology and same locality, we referred this new specimen to the same species and combined their scores into a single taxon as *Hanosaurus hupehensis* to maximize the morphologic information in other analyses here. Using the original matrix from Neenan et al.^16,19^, the basal position of *Hanosaurus* is recovered (Supplementary Fig. 4). Using the new matrix, our phylogenetic analyses conduct both maximum parsimony and Bayesian methods when the latter is employed to Sauropterygia for the first time (Materials and Methods). The topology generated by the consensus of the most parsimony trees (Supplementary Fig. 6-8) is fairly comparable to the Bayesian results (Supplementary Fig. 9, 10). Our phylogenetic results further confirm a monophyletic clade including all the eosauropterygians, placodontiforms, saurosphargids, and relevant Triassic marine reptiles (e.g., *Helveticosaurus* and *Atopodentatus*). Here we name this new and previously unnamed monophyletic clade as Sauropterygiformes (Fig. 2) and *Hanosaurus* is stably recognized as the basal-most taxon of this clade in all most-parsimony trees and Bayesian results.

**Figure 2.**
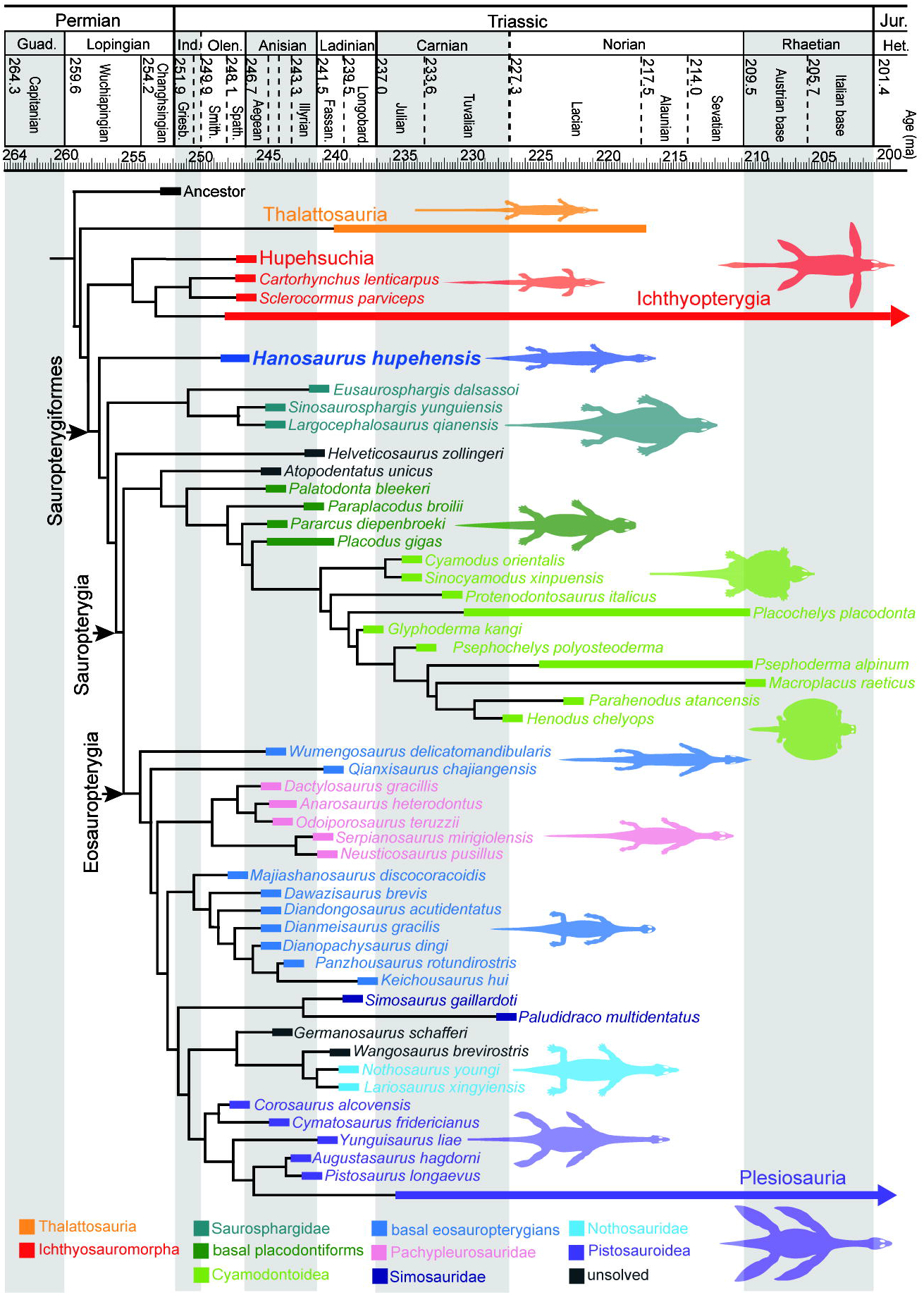
Dated cladogram of Sauropterygiformes based on the analyses of our new dataset on Triassic marine reptiles. Taxonomic groups are marked by different colors. Topologies are combined with results from both maximum parsimony and model likelihood (Supplementary Fig. 5-10). Colored bars of each taxon indicate the duration of discovered fossil remains. The ages of the nodes are obtained from the posterior median evaluations (Supplementary Fig. 11). Geologic time follows the latest monographs (Materials and Methods).

## DISCUSSION

### *Hanosaurus* as the basal-most sauropterygiform

Considering all the known sauropterygiform taxa on the genus level based on representative species for the first time, our work illustrates a more explicit phylogeny of sauropterygiform marine reptiles (Fig. 2, Supplementary Fig. 11). According to our analyses (see SI for more synapomorphies of main clades), *Hanosaurus* is recognized as a sauropterygiform reptile based on the possession of: an akinetic palate, a pectoral fenestration, gastralia segments with five elements, and the crown of marginal teeth with the concave lingual surface. Moreover, *Hanosaurus* is suggested to be the basal-most taxon of Sauropterygiformes in terms of the absence of: elongate transverse processes of dorsal vertebrae, radius longer than ulna, and curved humerus that all are shared by relatively less-basal sauropterygiforms. Additionally, *Hanosaurus* exhibits mosaic anatomical morphologies resembling saurosphargids, basal placodontiforms, and basal eosauropterygians. The general body size and shape, the cranial configuration, and the pachyostosis of the vertebral column, collectively reflect its close affinity to primitive eosauropterygians including the pachypleurosaur-like forms. However, compared to any of the known eosauropterygians, the neck of *Hanosaurus* is shorter and possesses fewer cervical vertebrae. Given the phylogenetic position of *Hanosaurus* as an early-diverging clade, it suggests that a neck of modest length and a cervical count of about 15 are primitive for Sauropterygiformes, and the neck became even shorter in Saurosphargidae and Placodontiformes when its length largely increased in Eosauropterygia, especially in the subgroup, Plesiosauria. Significantly, the disc-like coracoid and pubis, and less constricted ischium in *Hanosaurus* are reminiscent of saurosphargids, basal placodontiforms, and even earlier deviated thalattosaurians^20^ and hupehsuchians^15^. Nevertheless, differing from saurosphargids and placodontiforms, *Hanosaurus* also remains a primitive condition, lacking dermal armor, uncinate process on ribs, and further specialized teeth. The basal status of *Hanosaurus* and the phylogenetic result correspond to the fact that it represents one of the oldest members of Sauropterygiformes that occurred in the Early Triassic^21^.

### Body plan of early sauropterygiforms

Although as the basal-most sauropterygiform, *Hanosaurus* does not represent the terrestrial ancestor of sauropterygians. Apart from being preserved in marine sediments, *Hanosaurus* is certainly an aquatic animal based on its pachyostotic vertebral column achieving neutral buoyancy in the water, and the reduced ilium with bar-shaped sacral ribs that could not support the bodyweight to walk on land^22^. Besides these modifications, the slender and elongate trunk with the remarkably shortened limbs of *Hanosaurus* also featured aquatic adaptations as seen in other marine reptiles^23^. We illustrate the body shape of marine reptiles by direct measurements and ratios of various body regions (see Materials and Methods, and Supplementary Information). Terrestrial reptiles (e.g., *Araeoscelis* and *Vadasaurus*) normally have modest trunk lengths with considerable sizes of limbs (Fig. 3) for support and movement on land. Conversely, at the early stage of aquatic adaption in many marine reptilian groups (e.g., *Chaohusaurus*, an ichthypterygian; *Nanchangosaurus*, a hupehsuchian; *Pleurosaurus*, a marine rhynchocephalian), a body form combining an elongate trunk and shortened limbs is consistently present (Fig. 3). The elongate trunk can be additionally reflected by the higher number of dorsal vertebrae (Supplementary Fig. 12). Many marine reptiles also have a very long tail, but this long tail seems an ancestral feature inherited from their terrestrial ancestors^24^ like the early diapsids (Supplementary Fig. 12).

**Figure 3.**
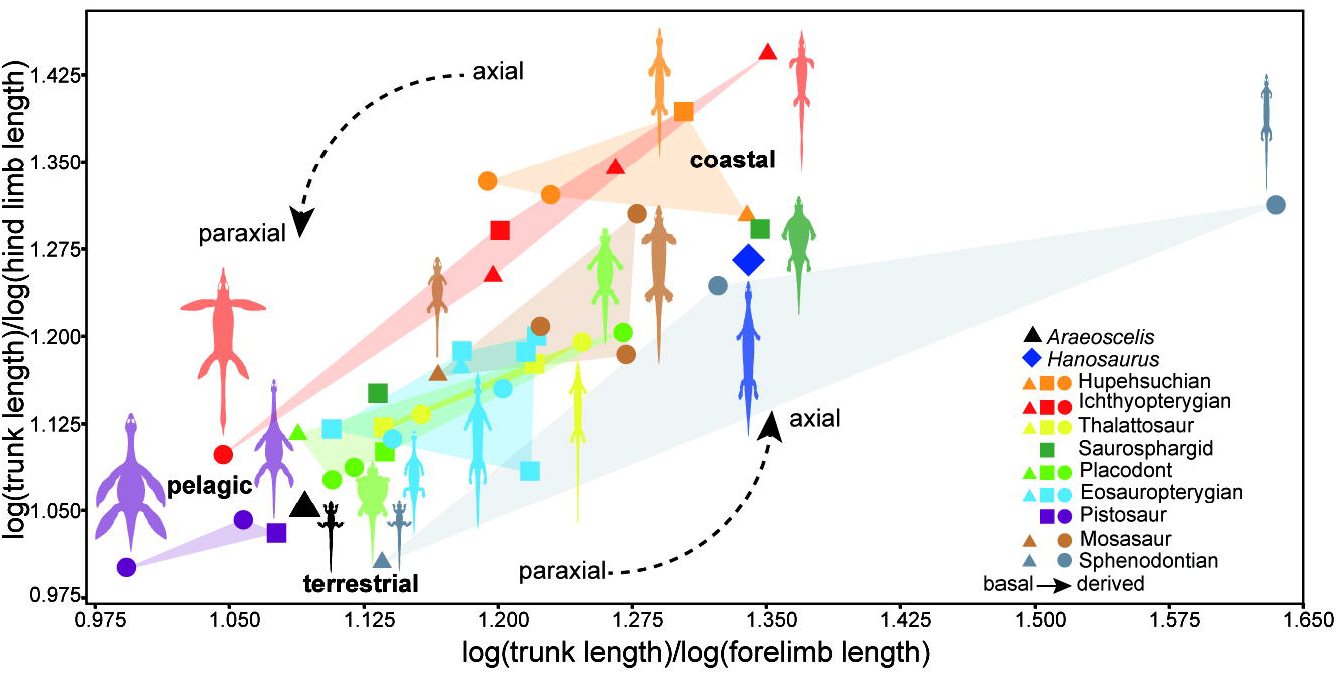
Convergence of body plan in secondarily aquatic reptiles reflected by the ratios of trunk/forelimb length and trunk/hind limb length. Taxonomic groups are labeled in different colors, and relative phylogenetic stages (basal, medium, derived) are plotted using varied shapes. Terrestrial, coastal marine, and pelagic marine taxa are respectively clustered in distinct areas. From basal to derived members, Sauropterygiforms and ichthyopterygians show a trend from axial to paraxial locomotion, while marine sphenodontians and mosasaurs are from paraxial to axial. This diagram illustrates the body plan convergence of long trunk and short limbs at the early aquatic adaptation stage.

In multiple lineages of marine reptiles, this ‘long-trunk’ convergence reflects the evolutionary constraints inheriting the axial undulatory movement^25^ from their terrestrial ancestors, and many basal members are anguilliform swimmers such as the primitive ichthyosauriforms^26,27^. Sauropterygia used to be considered as an exception^23^ because they were known as paraxial swimmers having relatively short and rigid trunks^28^ and using the limbs as thrusters, in which nothosaurs^29^ and plesiosaurs are representatives. However, *Hanosaurus* possesses more dorsal vertebrae (Supplementary Fig. 12) and evidently shortened fore- and hind-limbs compared to other sauropterygiforms, but more similar to other aquatic reptiles like *Nachangosaurus* and *Chaohusaurus* in this aspect (Fig. 3, Supplementary Fig. 12, 13). With the discovery of *Hanosaurus*, we herein propose that the basal sauropterygiform marine reptiles are also dominated by the evolutionary constraint to evolve an elongate trunk and short limbs at their early stage of secondarily aquatic adaptation. This ‘long-trunk’ convergent in marine reptiles (Fig. 3, Supplementary Fig. 14) is probably selected by the coastal marine environment full of rocks and reefs, and/or caused by the mutation of dorsal vertebra duplication^30^ that is possibly easier to occur compared to the podial modifications in limbs. Although the pachyostotic dorsal ribs and the dense gastral basket preclude flexion of the trunk, the marine reptiles developing pachyostosis could be anguilliform swimmers^31^. We speculate that *Hanosaurus* and early sauropterygiform reptiles have axial undulatory locomotion at least more than most of sauropterygians if not as much as that in ichthyosauriforms. Therefore, before evolving the underwater-flying stroke of pistosaurs, the elongate neck in eosauropterygians, and the turtle-like appearance in placodonts and saurosphargids (Fig. 2), *Hanosaurus* shows an unexpected ‘long-trunk’ body plan as a possible anguilliform swimmer at the beginning of sauropterygiform evolution.

### Adaptive radiation of early sauropterygiforms

After the ‘long-trunk’ convergent evolution of the basal-most members at the very early stage, sauropterygiforms show a divergent evolution to generate Saurosphargidae with laterally expanded flat trunks, Placodontia with a body covered with heavy armours superficially resembling turtles, and Eosauropterygia including abundant pachypleurosaur-like small members and large plesiosaurs developing long necks and broad fins (Fig. 2). These different sauropterygiform clades evolved manifold phenotypes occupying a large region in the morphospace (Fig. 4, Supplementary Fig. 13). In comparison, ichthyopterygians have less change from anguilliform *Chaohusaurus* to dolphin-like derived ichthyosaurs, and thalattosaurians show little bauplan modification besides the changes of head shape and neck length (Fig. 4). Based on our latest dataset and phylogenetic results of Sauropterygiformes, our tip-dating result indicates that Sauropterygiform marine reptiles radiated rapidly (Supplementary Fig. 11) from the latest Permian to the earliest Triassic (Fig. 2). High rates are estimated at the beginning of sauropterygiform evolution, and the three main subclades (Saurosphargidae, Placodontiformes, and Eosauropterygia) arose within less than 10Ma from the latest Permian. When more taxa of early sauropterygiforms were added, the high evolution rates among placodonts^32^ are not recovered. Another episode of rapid evolution is at the early stage of eusauropterygian clades leading to the large-sized nothosaurs and pistosaurs (Supplementary Fig. 11). Apart from the taxon diversity with more than 40 genera discovered, under our updated phylogenetic framework, sauropterygiform reptiles took full advantage of the newly emerged marine ecological opportunities in its early evolutionary burst by evolving novel morphologies with associated locomotion^33^, habitats, and diets. Sauropterygiforms soon rose from the aftermath of the end-Permian mass extinction to become a predominant reptilian group conquering the sea.

**Figure 4.**
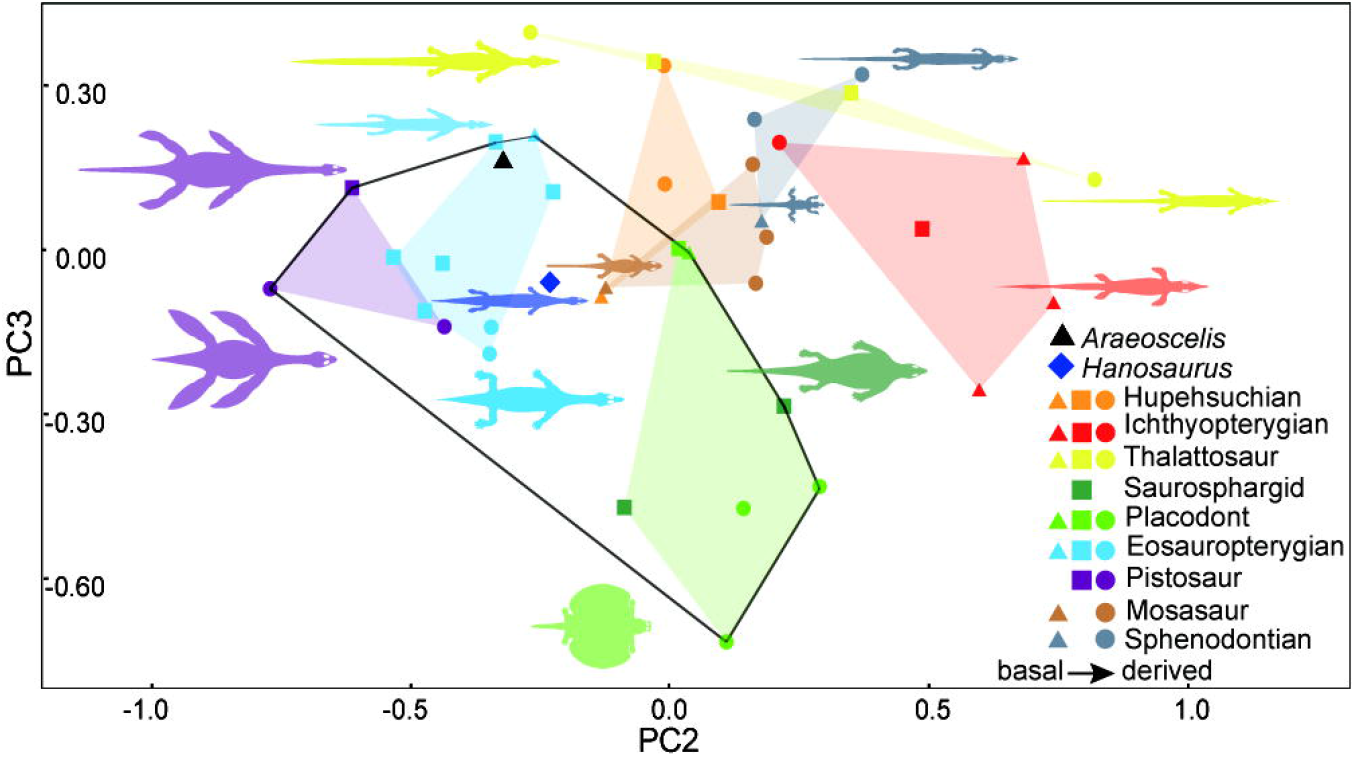
Morphospace based on principal component (PC)2 and PC3. Body shape principal component analysis reflects the morphologic variances among different Mesozoic marine reptiles. Black lines outline the area occupied by sauropterygiform reptiles.

## STAR METHODS

### Fossil materials

The new specimen (IVPP V 15911) and the holotype (IVPP V 3231) of *Hanosaurus hupehensis* are permanently deposited at the Institute of Vertebrate Paleontology and Paleoanthropology (IVPP), Beijing, China. As for other observed specimens, we had examined the specimens as many as possible, especially in the richest collections of Triassic marine reptiles in China and European countries. We (at least one of the authors) examined the true specimens of more than 80% of the total sampled taxa (Supplementary Table. 1). The leading author (WW) observed more than 70%. One or more of the authors conducted a detailed study on about 50% of the total taxa. To avoid bias due to the observation and interpretations of different authors, the state codes for all characters were carefully scored by the leading author (WW) with a consultation with other authors. As for the genera including several species, if possible, we chose a species that is represented by complete, well-preserved adult specimens (Supplementary Table. 1).

### Character matrix construction

To investigate the phylogenetic relationships of *Hanosaurus* among marine reptiles, we constructed an updated morphological dataset (62 taxa, 181 characters) focusing on Triassic sauropterygians and relevant reptiles in terms of a series of previous character matrix and character state coding were largely based on our firsthand observation of specimens. Compared to the previous most comprehensive dataset of sauropterygians^16,17^, we added 26 new taxa to include all the known placodontiforms, eosauropterygians, and saurosphargids as the ingroups or the target groups herein, and sampled the taxa more adequately and representatively than any of the previous datasets^18^ to avoid involving redundant species of one genus^34^. *Youngina capensis* that could represent the characters of primitive diapsids was used to root the cladogram. According to the latest studies from multiple independent research groups^16,17,32,35,36^ and our phylogenetic results using the previous matrix (Supplementary Fig. 3, 4), Ichthyopterygia and Thalattosauria have been frequently designated as two successive sister groups of Sauropterygia. Therefore, with more primitive members of Ichthyosauriformes^3,37^ discovered from eastern Tethyan fauna, instead of using the whole group of Ichthyopterygia and Thalattosauria as operational taxonomic units in the previous studies^16,17,19^, we used specific species of the basal-most and well-documented members of ichthyosauriforms (hupehsuchians included) and thalattosaurians as the outgroups to investigate the relationship among sauropterygian and saurosphargid taxa. Although the monophyly of the marine reptile group combining Ichthyopterygia, Thalattosauria, and Sauropterygia is controversial, it is beyond our scope here focusing on the inner relationship within sauropterygians and their relationship with unsolved taxa such as saurosphargids and *Atopodentatus*. A larger matrix involving the basal members of Archosauromorpha and Lepidosauromorpha will be considered in our future work when their inner relationships^35,36,38,39^ are better resolved respectively and more specimens are personally checked by us. The taxa of stem turtles (or pantestudines) were possibly related to sauropterygians in a few recent studies^40,41^, and we first-hand observed some specimens^41,42^ and tentatively added them in our matrix to double test. We added 40 new characters according to the homology principle and excluded the features dominantly affected by their aquatic lifestyle. We scored the character states based on the best-preserved adult specimens to decrease the ontogenetic, sexual, and individual variations. Under these criteria, our character matrix can be considered on the genus level, the species level, or even the specimen level, being available when extra operational taxonomic units on any level are added in a future study.

### Phylogenetic analyses and tip dating

To obtain the phylogeny of sauropterygian and allied marine reptiles such as saurosphargids, we conducted the maximum parsimony inference using TNT version 1.5^43^. All the characters were equally weighted and unordered. The general RAM in the memory was adjusted to 500 MBytes. Because the number of taxa was fewer than 100 and new technology search was unnecessary, herein we conducted the traditional search with random additional sequence and tree bisection-reconnection (TBR). The heuristic search ran 1000 replications of Wagner trees with one random seed and 10 trees saved per run. We conducted six rounds of maximum parsimony analyses. In the first round of analyses, we employed the data matrix of Neenan and colleagues^16,17,19^, and coded the character states of the holotype (IVPP V 3231) and the new specimen (IVPP V 15911) separately, when more informative scores from the skull were obtained in the holotype but more from the postcranial skeleton were coded in the new specimen respectively (Supplementary Data 1, used with the operational taxonomic unit (OTU) *Hanosaurus hupehensis* excluded). In the second round, considering the result (Supplementary Fig. 3) when two specimens were coded separately in the previous data matrix, we combined the information into one OTU as *Hanosaurus hupehensi*s (Supplementary Data 1, used when V 3231 and V 15911 excluded; Supplementary Fig. 4). In the third round, we coded IVPP V 3231 and IVPP V 15911 separately again in our updated data matrix (Supplementary Data 2, used with the OTU *Hanosaurus hupehensis* excluded) to further test the affiliation of these two specimens (Supplementary Fig., 5). In the fourth round, the scores were combined as one OTU *Hanosaurus hupehensis* (Supplementary Data 2, used when V 3231 and V 15911 excluded) with all 62 taxa involved (Supplementary Fig. 6). Based on the result of the fourth round, we conducted the reduced consensus tree analysis to find the wildcard OTUs, which came to be the stem pantestudines and hupehsuchians. Therefore, we excluded the stem pantestudines in our fifth round (Supplementary Fig. 7), and removed both pantestudines and hupehsuchians in the sixth round of analyses (Supplementary Fig. 8).

To further examine the phylogeny of sauropterygians and saurosphargids, we employed Bayesian phylogenetic inference using MrBayes version 3.2.7^44^. Sauropterygians were all extinct without extant descendants, and our character matrix included only morphological data, therefore, we used the Mkv model^45^ with variable ascertainment bias, equal state frequencies, and gamma rate variation across characters and the prior for the gamma shape parameter was exponential (1.0). For the Markov chain Monte Carlo (MCMC), two independent runs and four chains, including a cold chain and three hot chains, were used per run for 10 million generations and sampled every 1000 generations. The average standard deviation of split frequencies (ASDF) was less than 0.01, and the effective sample size (ESS) was larger than 100 both indicating an adequate result. We conducted two rounds of Bayesian analyses when *Hanosaurus hupehensis* was scored based on both two specimens. All 62 OTUs (Supplementary Data 3) were involved in the first round (Supplementary Fig. 9), and 54 OTUs with pantestudines and hupehsuchians removed in the second round (Supplementary Fig. 10).

To investigate the evolutionary rates and the divergence times among all sauropterygiform clades, we performed a Bayesian tip-dating (Supplementary Fig. 11) using the new data matrix with an unpartitioned model and the constrained parsimony tree topology in Mrbayes.3.2.7^44^. We used an independent lognormal^46^ relaxed clock model and the Mkv model^45^ with a gamma rate variation^47^ across all characters for likelihood calculation. The parameters of the MCMC process were the same as the above undated Bayesian phylogenetic analyses. The time tree was assigned a uniform prior^48^, and the root age had an offset exponential prior with a mean of 256.3 Ma and a minimum of 251.9 Ma. The ages of all marine reptile genera/species in the matrix were assigned uniform priors respectively with their lower and upper bounds of the corresponding geologic time stages^49^ and stratigraphic ranges (Supplementary Table. 2), except for *Psephoderma alpinum* being fixed at its last appearance date (201.4 Ma) because of having the youngest last appearance date among all taxa we considered.

### Specimen measurements and body shape analyses

We assembled a series of linear measurements from representative marine reptiles (Supplementary Data. 4). Our dataset includes all sauropterygiform subgroups and other major lineages of Mesozoic marine reptiles such as Hupehsuchia, Ichthyopterygia, Thalattosauria, marine lepidosaurs including marine sphenodontians and mosasaurs. We sampled and measured 40 nearly complete skeletons of 40 species/genera representing basal to crown taxa of each group (Supplementary Data. 4). According to our phylogenetic results and literature, we reconstructed a cladogram of all these sampled marine reptilian taxa (Supplementary Fig. 12). Measurements of the dataset were made either using a vernier caliper and flexible rule directly from the specimens or using the ImageJ program from the published photos and corresponding scale bars in literature. Our dataset included nine discrete and unrelated linear values (Supplementary Data. 4): the skull length (SL) from the anterior margin of the snout to the posterior end of the occiput, the skull width (SW) as the maximum width of the dermal cranium, the neck length (NL), the body length (BL) (same as the trunk length in this study) from the first dorsal to the last sacral vertebra, the body width (BW) (same as the trunk width in this study) measured from the maximum width of the rib cage when the specimen was preserved in dorsal view or the length of the longest rib in lateral view, the tail length (TL), the tail width (TW) as the maximum width of the proximal region of the tail in dorsal view or the width of the largest caudal centrum including the length of the associated transverse process, the forelimb length (FL), and the hind limb length (HL).

Our analyses were conducted using the paleontological statistics software package (PAST) version 4.0^50^. To test the convergence of long trunk and short limbs, we first conjunctively considered these two features and made a binary plot using the dimensionless ratios of BL/FL and BL/HL as the coordinates with all original measurements (BL, Fl, and HL) already log_10_-transformed (Fig. 3). Secondly, we made a bar graph based on the ratio of BL/BW to show the elongation of the trunk itself (Supplementary Fig. 12). Thirdly, we created a stacked bar chart of NL/BL/TL to compare the percentages of these three parts in different taxa (Supplementary Fig. 12). These results should be comprehensively considered, because some features might be ignored due to others, such as that the BL of *Henodus* (a highly specialized placodont) occupied a large percentage in the entire length but its trunk was not actually elongated, while the body of *Pleurosaurus* (a marine sphenodontian) was extremely long but not that notable with a tail even longer. These nine measurements could outline the body shapes, therefore, we collectively used aforementioned nine linear measurements being log_10_-transformed and employed principal component analysis (PCA) to demonstrate the shape variances of the sampled marine reptiles. We also conducted principal coordinate analysis (PCoA), which showed almost the same results as PCA herein. The principal component 1 axis (PC1) reflected about 86% of the variance showing positive eigenvector coefficients with all the nine measurements, hence PC1 seemed mainly related to the size with large marine reptiles on one side and small ones on the other (Supplementary Fig. 13). PC2 described more than 7% of the variance being evidently negative (−81%) with the neck length, which explained the elongation or shorten of the cervical region with *Plesiosaurus* and *Cartorhynchus* (an ichthyosauriform) at the two opposite ends. PC3 explained over 3% of the variance, and its eigenvector coefficients were mostly negative (−64%) with the body width and mostly positive (61%) with the tail length when *Henodus* and *Anshunsaurus* (a thalattosaurian) were at the two extremes respectively. We preferred the binary morphospace using PC2 and PC3 to demonstrate the shape disparities of multiple marine reptilian lineages (Fig. 4).

## Supporting information

Supplementary Information_Ancestral Body Plan and Adaptive Radiation of Sauropterygian Marine Reptiles_Wang et al. preprint

Supplemental Data 1. Neenan Matrix Add Hanosaurus V3231 V15911 for TNT_preprint

Supplemental Data 2. Triassic marine diapsid matrix for TNT

Supplemental Data 3. Triassic marine diapsid matrix for Bayes

Supplemental Data 4. Measurements of representative marine reptiles

## SUPPLEMENTAL INFORMATION

is available for this paper.

## ACKNOWLEDGMENTS

We thank J. Z. Ding for specimen preparation; W. Gao for photographing; G. Ugueto for fauna reconstruction; C. Zhang, Y. L. Yu, and H. B. Wang for assistance on Bayesian analyses; Z. H. Zhou, X. Xu, and X. J. Ni for discussion; R. X. Zhu, C. L. Deng, and H. Y. He for support. W. W. thanks T. M. Scheyer, O. W. M. Rauhut, E. E. Maxwell, R. R. Schoch, I. Werneburg, S. Nosotti, L. J. Zhao, and D. Y. Jiang for their hospitality and access to the collections. The study was supported by the Strategic Priority Research Program (B) of the Chinese Academy of Sciences (XDB26000000), the Youth Innovation Promotion Association of Chinese Academy of Sciences, the National Natural Science Foundation of China (42002019 and 41972014), State Key Laboratory of Paleobiology and Stratigraphy (Nanjing Institute of Geology and Palaeontology, CAS) (20203119), Key Laboratory of Vertebrate Evolution and Human Origins of Chinese Academy of Sciences (IVPP, CAS) (LVEH020001).

## AUTHOR CONTRIBUTIONS

C. L. and W. W. conceived the research; C. L., Q. S., L. C., and W. W. participated the fieldwork; W. W. undertook the morphological comparisons, data collection, figure preparation, phylogenetic and statistical analyses. W. W., X. W., and C. L. interpreted the fossil and wrote the manuscript with feedback from Q. S. and L. C.

