## Supplementary Information_Ancestral Body Plan and Adaptive Radiation of Sauropterygian Marine Reptiles_Wang et al. preprint for "Ancestral Body Plan and Adaptive Radiation of Sauropterygian Marine Reptiles"

Contents of Supplementary Information

1. Additional description and comparison
2. Character list for phylogenetic analyses
3. Specimen list of the sampled Triassic marine diapsids
4. Phylogenetic results and clade synapomorphies
5. Taxon ages of Triassic marine reptiles
6. Measurements of secondarily aquatic reptiles

### 1. Additional description and comparison

Abbreviations in Fig. 1: an, angular; ar, articular; as, astragalus; ax, axis; cal, calcaneum; car, caudal rib; cav, caudal vertebra; cl, clavicle; cn, coronoid; co, coracoid; cp r, capitulum of rib; cr, cervical rib; cv, cervical vertebra; d, dentary; dc, distal carpal; dp, diapophysis; dr, dorsal rib; dt, distal tarsal; fe, femur; fi, fibula; ga, gastralia; h, humerus; hy, hyobranchium; icl, interclavicle; il, ilium; im, intermedium; ipv, interpterygoid vacuity; is, ischium; lvr, lateroventral ridge; m, maxilla; mc, metacarpal; mt, metatarsal; ob, orbit; oc, occipital condyle; pl, palatine; pp, parapophysis; prz, prezygapophysis; pt, pterygoid; ptf, pterygoid flange; pu, pubis; q, quadrate; ra, radius; sa, surangular; sc, scapula; sp, splenial; stf, supratemporal fenestra; sv, sacral vertebra; tb r, tuberculum of rib; ti, tibia; tp, transverse process; ul, ulna; uln, ulnare.

Here we describe more morphology in detail of the new specimen, IVPP V 15911. Preserved in a single slab of limestone, it is a nearly complete skeleton that lacks only the posterior caudal vertebrae and a few phalanges (Fig. 1 a, b). The complete skull is preserved in ventral view and it is dorso-ventrally compressed. The crack puts the rostral part on a level higher than that of the rest skull (Fig.1 c, d). Anterior to the thin, reduced, and longitudinally oriented pterygoid flanges, an array of tiny papillae are present on the pterygoid, but whether these are alveoli is uncertain. The dentition is incompletely preserved and partially exposed in the ventral view. There is a crack separating the anterior rostral part from the lateral cheek, and the suture between the premaxilla and the maxilla is uncertain. Therefore, the accurate numbers of premaxillary, maxillary, and dentary teeth respectively are unknown. According to the positions and orientations, anterior to the crack, we can identify 6 teeth on the right dentary, 5 on the left dentary, and 2 on the anterior portion of the left maxilla. About 5 premaxillary teeth are estimated in accordance with the anterior teeth on the dentary. Posterior to the crack, lateral teeth are exposed only on the right side to be interpreted as 8 maxillary teeth and 3 dentary teeth at least. From the occipital-cervical articulation, 15 cervical vertebrae are identified including the atlas and the axis. We identify the first dorsal rib based on the articulation to the sternum instead of the position anterior to the pectoral girdle. The 15th cervical rib resembles but is more slender and shorter than the first dorsal rib, and its distal end is not expanded as in the first dorsal rib (Fig.1 a, b, g, h). The centrum of cervical vertebrae 2 to 13 is stout and possesses a pair of longitudinal ridges along its ventrolateral surface, which are identified as the parapophysis to articulate to the capitulum of the cervical rib, and the ventral surface between the ridges is smooth and longitudinally shallowly concave. The parapophysis is absent in the last one or two cervical, all the dorsal, sacral, and caudal centra that are constricted and also show a concave ventral surface. The posterior caudal centra gradually decrease in size but are still stout. The anterior cervical ribs are bicephalous and develop free anterior processes, and the dorsal and caudal ribs are holocephalous (Fig. 1 e, f; Supplementary Fig. 2 i, j, m, n). The chevron is short and simply v-shaped. The gastralia are well developed between the pectoral girdle and the pelvic girdle. There are more than 70 sets of gastralia closely arranged but not overlapped to one another, and each gastralium set is composed of five flattened segments (Fig. 1 g, h). The pectoral girdle (Fig. 1 g, h; Supplementary Fig. 2 a, b) is completely preserved and exposed in ventral view, resembling the typical interclavicle-clavicle articulation pattern of sauropterygian but the clavicle is more anterior to the scapula than sauropterygians. Although the interclavicle is incomplete due to erosion, it seems short and rhomboidal without the posterior process. The anterior end of the clavicle closely approaches one another but does not meet in ventral view. The ventral portion of the scapula is robust developing a large articular surface to the anterolateral margin of the coracoid. The base of the scapula blade can be identified according to its limited exposure in between the humeral proximal head and the rib. Both forelimbs are nearly complete only with a few carpals and digital elements missing (Fig. 1 g, h). The radius is slightly curved, with no proximal or distal expansions. The ulna is rather straight, with the shaft slightly constricted, and its proximal end is slightly wider than the distal one. The metacarpal I is stout and half the length of the metacarpal II. Metacarpal II and IV are similar in length, metacarpal III is slightly longer than the former two, and metacarpal V is slightly shorter than metacarpals II and IV. The phalangeal formula of the manus is possibly 2-3-4-5(?)-3(?) (Fig. 1 g, h; Supplementary Fig. 2 a, b). The pelvic girdle is nearly complete, with certain parts damaged by erosion. The right pubis and the left ischium are perfectly preserved and exposed in ventral view (Fig. 1 i, j). The ilium is rather reduced but still with an evident anterior process. A pair of the hind limbs are exposed in ventral view, with the left one better preserved. The distal femoral condyles are weakly developed, with a smooth surface. Metatarsal I is obviously shorter than the others, metatarsal III is much longer than the others, and metatarsals II, IV, and V are similar in length and width. The phalangeal formula of the pes is 2-3(?)-4-5-3 (Fig. 1 i, j).

We consider that two specimens of *Hanosaurus* reached their adulthood size compared with many pachypleurosaur or similar taxa. The preserved length of the entire new skeleton is about 80cm, and the estimated total length of this animal may have approximately reached 1 meter. After the comparison of the femur, IVPP V 15911 (about 33 mm long) is about 75% of the length of IVPP V 3231 (about 44 mm long), implying that the latter may have reached a length of 125 cm. *Hanosaurus* is a mediumsized sauropterygiform and is comparable in length to the basal eosauropterygians *Wumengosaurus* and *Qianxisaurus*. The common size of most European pachypleurosaurs and Chinese pachypleurosaur-like eosauropterygians (e.g., *Dawazisaurus*, *Diandongosaurus*, and *Keichousaurus*) is around 50 cm in length, which is much smaller compared with the members of the later clades such as Nothosauroidea. Moreover, the adulthood of these two specimens of *Hanosaurus* is supported by some anatomical features, such as the well-developed long bones of all four limbs, well-demarcated marginal edges and prominent muscle scars on the femurs, and the fully ossified carpals and tarsals (Fig. 1 g, h, i, j; Supplementary Fig. 2 m, n).

The new specimen (IVPP V 15911) and the type specimen (IVPP V 3231) came from the same horizon showing the same lithology of the very neighboring locality in the middle-west region in Hubei province, China (Supplementary Fig. 14). After our further examination of the type specimen, we confirm more synapomorphies to support that our new specimen can be referred to *Hanosaurus hupehensis*. The two specimens are exclusively similar in these features: the maxillary or posterior teeth possess a constricted base and a crown with a concave lingual surface (Supplementary Fig. 1, 2 c, d, e, f); the retroarticular process is distinctly large (Fig. 1 c, d; Supplementary Fig. 1); the cervical ribs process a prominent free anterior process (Supplementary Fig. 2 i, j, k, l); the pubis and ischium are similarly roundish and kidney-shaped, respectively; and the tarsals are the same in the number of ossifications (Supplementary Fig. 2 m, n, o, p). More morphological characters are documented in our data matrix when IVPP V 3231 and IVPP V 15911 were respectively coded.

Notably, from the same horizon and adjacent localities, other three specimens, YIGM V 0940, YIGM V 0941, and HFUT YZS-16-01, were reported and identified as *Lariosaurus sanxiaensis*, distinguished from *Hanosaurus hupehensis* currently only by the count of ossified tarsals and the shape of coracoid according to the descriptions in previous studies, when the tarsal number is usually varied in different individuals or ontogenetic stages and the coracoid is poorly preserved in IVPP V 3231. However, in HFUT YZS-16-01, the exact same morphology of the coracoid, the clavicle, the dorsal ribs, and the femur (possibly misinterpreted by Li and Liu as 'ulna') can be confirmed in the type specimen of *Hanosaurus hupehensis* and this new skeleton. Additionally, these three specimens show clear different characters from all taxa of Nothosauridae, the monophyletic family including *Nothosaurus* and *Lariosaurus* species. On the contrary from of aforementioned three specimens, nothosaurid species exhibit the supratemporal fenestra much larger than the orbit, an evidently larger amount of the cervical vertebrae, an obvious constriction at the mid-point of the coracoid and pubis, the ulna much broader than the radius, the humerus shorter than the femur and so on. Here we tentatively refer HFUT YZS-16-01 to *Hanosaurus hupehensis*, and we temporarily prefer not to consider *Lariosaurus sanxiaensis* in our analyses and discussion because of the current poor understanding of this taxon. In our separate future studies, more detailed comparisons and more unprepared fossil material will be involved to clarify the differences and the validity between these two taxa.

#### 2. Character list for phylogenetic analyses

We construct a new character list for our data matrix involving 181 characters. Of which 141 characters are from the latest studies on Sauropterygia (Neenan et al., 2013) and related taxa (Scheyer et al., 2017). The other 40 characters (bold font) include those newly added by us as well as those cited/modified from the specific studies on Placodontiformes (Wang et al., 2019), Pachypleurosauria (Rieppel and Lin, 1995), Nothosauria (Liu et al., 2014), Thalattosauria (Liu et al., 2013), and Ichthyopterygia (Ji et al., 2016). The sequence of characters corresponds, on the one hand, to those of Neenan et al. (2013) for easy comparison, and, on the other hand, follows the manner of anatomical observation.

(1) Bones in dermatocranium: distinctly sculptured (0); relatively smooth (1).

(2) Preorbital and postorbital region of skull: of subequal length (0); preorbital region distinctly longer (1); postorbital region distinctly longer (2).

(3) Snout: relatively short (0); elongated with broad anterior termination (1); elongated and tapering anteriorly (2); elongated and spatulate (3) (Wang et al., 2019: 3).

(4) Distinct snout constriction in adult: absent (0); present (1).

(5) Premaxillae: small (0); large, forming most of snout in front of external nares (1).

**(6) Anterior portion of premaxilla: nearly horizontal (0); turned downwards (1). (modified from Liu et al., 2013: 1)**

(7) Postnarial process of premaxilla: absent (0); present, excluding maxilla from posterior margin of external naris (1).

**(8) The ventral surface of the premaxilla is level with the ventral surface of the maxilla (0) or the premaxilla is distinctly downturned (1). (Wang et al., 2019: 4)**

(9) External nares: not retracted (0); retracted with a longitudinal diameter approaching or exceeding half the longitudinal diameter of orbit (1); retracted, narrow, and with a longitudinal diameter distinctly less than half the longitudinal diameter of orbit (2).

(10) Nasal(s): shorter than frontal(s) (0); longer than frontal(s) (1).

(11) Nasal(s): not reduced (0); reduced (1); absent (2).

**(12) Nasal anteriorly extending beyond external naris: false (0); true (1). (Ji et al., 2016: 10)**

(13) Nasal(s): meeting in dorsomedial suture (0); fused (1); separated from one another by nasal processes of premaxillae extending back to frontal(s) and/or anterior processes of the frontals (2).

**(14) Nasal-prefrontal contact: present (0); absent (1). (Liu et al., 2014: 14)**

**(15) Nasal: does not extend posterior to level of anterior margin of orbit (0); does extend posteriorly beyond this level (1). (Liu et al., 2013: 9)**

(16) Lacrimal: present, entering external naris (0); present, excluded from external naris (1); absent (2).

**(17) Maxilla-prefrontal contact: absent (0); present (1). (Ji et al., 2016: 5)**

(18) Dorsal exposure of prefrontal: large (0); reduced (1).

(19) Prefrontal: without slender anteromedial process (0); with slender anteromedial process entering between maxilla and premaxilla (1).

**(20) Frontal: broadly enters the dorsal margin of the orbit (0); restrictedly enters due to an elongated anterior process of the postfrontal (1); restrictedly enters due to an elongated posterior process of the prefrontal (2); remains excluded from the dorsal margin of the orbit due to a prefrontal-postfrontal contact (3). (modified from Rieppel and Lin, 1995)**

(21) Frontal(s) in adult: paired (0); fused (1).

**(22) Anterolateral processes of frontals well developed (0) or reduced (1). (Wang et al., 2019: 11)**

**(23) Parietal without (0) or with (1) distinct anterolateral processes embraced by postfrontal and frontal. (Wang et al., 2019: 12)**

(24) Distinct posterolateral processes of frontal(s): (0) absent; (1) present.

(25) Frontal: widely separated from upper temporal fossa (0); narrowly approaching upper temporal fossa (1); entering the anteromedial margin of upper temporal fossa (2).

**(26) Frontals do not (0) or do (1) reach posteriorly beyond the level of the anterior margin of the upper temporal fossa. (Wang et al., 2019: 13)**

(27) Postfrontal: large and plate-like (0); with distinct lateral process overlapping the dorsal tip of postorbital (1); with reduced lateral process and hence more of an elongate shape (2).

**(28) Postfrontal and postorbital: separate (0); fused (1). (Liu et al., 2013: 15)**

(29) Jugal: extending anteriorly along the ventral margin of orbit (0); restricted to a position behind orbit but entering the latter's posterior margin (1); restricted to a position behind orbit without reaching the latter's posterior margin (2).

(30) Jugal: extending backward no farther than to the middle of cheek region (0); extending nearly to the posterior end of skull (1).

(31) Jugal: excluded from upper temporal arch (0); entering upper temporal arch (1).

**(32) Jugal–squamosal contact along dorsal/proximal margin of supratemporal fenestra: absent (0), or present (1). (Wang et al., 2019: 56)**

(33) Parietal(s) in adult: paired (0); fused in their posterior part only (1); fully fused (2).

(34) Parietal skull table: broad (0); weakly constricted (1); strongly constricted (at least posteriorly) (2); forming a sagittal crest (3).

(35) Pineal foramen: close to the middle of skull table (0); weakly displaced posteriorly (1); strongly displaced posteriorly (2); displaced anteriorly (3); absent (4).

**(36) Pineal foramen located within a deep trough: absent (0); present (1). (Liu et al., 2014: 16)**

**(37) Postorbital: largely forms the anterolateral margin of upper temporal fossa (0); restrictedly enters the anterolateral margin of upper temporal fossa (1). (New character)**

(38) Postparietals: present (0); absent (1).

(39) Tabulars: present (0); absent (1).

(40) Supratemporals: present (0); absent (1).

(41) Temporal region of skull: relatively high (0); strongly depressed (1).

(42) Upper temporal fossa: absent (0); present and subequal in size or slightly larger than orbit (1); present and distinctly larger than orbit (2); present and distinctly smaller than orbit (3); secondarily closed (4).

(43) The anteromedial corner of upper temporal fossa: not (0); partially (1); fully (2) floored by a descensus from postorbital, which together with neighboring elements (postfrontal, parietal) separates it from orbit.

(44) Lower temporal fossa: absent (0); present and closed ventrally (1); present but open ventrally (2).

(45) Squamosal: descending to ventral margin of skull (0); broadly separated from ventral margin of skull (1).

(46) A box-like suspensorium of squamosal: absent (0); present (1).

(47) Distinct notch of squamosal to receive distal tip of paroccipital process: absent (0); present (1).

(48) Quadratojugal: present (0); absent (1).

(49) Anterior process of quadratojugal: present (0); absent (1).

(50) Quadrate: covered by squamosal and quadratojugal in lateral view (0); exposed in lateral view (1).

(51) Posterior margin of quadrate: straight (0); concave (1).

(52) Lateral conch on quadrate: absent (0); present (1).

**(53) Scleral ossicles: present (0) or absent (1). (Rieppel and Lin, 1995: 21)**

(54) Dorsal wing of epipterygoid: approximately as broad as its base (0); narrower than its base (1).

(55) Braincase: located at posterior end (0); deeply recessed below parietal skull roof (or parietal sagittal crest) (1).

(56) Occipital crest: absent (0); present but squamosals not meeting behind parietal (1); present and squamosals meeting behind parietal (2).

(57) Occiput: with paroccipital process forming the lower margin of posttemporal fossa and extending laterally (0); paroccipital processes trending posteriorly (1); plate-like with no distinct paroccipital process and with strongly reduced posttemporal fossae (2).

(58) Mandibular articulations: approximately at level with occipital condyle (0); displaced to a level distinctly behind occipital condyle (1); positioned anterior to occipital condyle (2).

(59) Supraoccipital: exposed more or less vertically on occiput (0); exposed more or less horizontally at posterior end of parietal skull table (1); U-shaped (2).

(60) Contact between exoccipitals and basioccipital condyle: present (0); absent (1).

(61) Basioccipital tubera: free (0); in complex relation to pterygoid, as they extend ventrally (1); in complex relation to pterygoid, as they extend laterally (2).

(62) Palate: kinetic (0); akinetic (1).

(63) Premaxillae: entering internal naris (0); excluded from internal naris (1).

(64) Posterior palatine vacuities: absent (0); present (1).

**(65) Palatines: separated, at least partially, by long pterygoids (0), or meeting in medial suture (1). (Wang et al., 2019: 60)**

(66) Pterygoids: longer than palatines (0); shorter than palatines (1).

(67) Pterygoid flanges: well-developed and transversely oriented (0); well developed and longitudinally oriented (1); strongly reduced (2).

(68) Ectopterygoid: present (0); absent (1).

(69) Suborbital fenestra: absent (0); present (1).

(70) Internal carotid passage: entering basicranium (0); entering quadrate ramus of pterygoid (1).

(71) Splenial bone: entering mandibular symphysis (0); excluded therefrom (1).

(72) Distinct coronoid process of lower jaw: absent (0); present (1).

(73) Strongly projecting lateral ridge of surangular defining the insertion area for superficial adductor muscle fibres on the lateral surface of lower jaw: absent (0); present (1).

(74) Mandibular symphysis: short (0); somewhat enforced (1); elongated and 'scoop-like' (2).

**(75) Dorsal margin of dentary symphysis: straight (0); recurved (1). (Liu et al., 2013: 21)**

**(76) Mandible constriction: absent (0); present (1). (New character)**

(77) Retroarticular process of lower jaw: absent (0); present (1).

(78) Trough on dorsal surface of retroarticular process: absent (0); present (1).

**(79) Dentigerous region in adult: complete (0); largely reduced (1); edentulous (2). (Ji et al., 2016: 57)**

(80) Teeth: setting in shallow or deep sockets (0); superficially attached to bone (1).

**(81) Overbite: (0) absent or very slightly; (1) present. (Ji et al., 2016: 50)**

(82) Durophagous dentition: absent (0); present (1).

**(83) Number of premaxillary teeth: four or more (0); three to one (1); modified into a single row of denticles (2); absent (3) (modified from Wang et al., 2019: 35).**

(84) Anterior (premaxillary and dentary) teeth: upright or only slightly procumbent (0); strongly procumbent (1); absent (2).

(85) Premaxillary and anterior dentary fangs: absent (0); present (1).

(86) One or two enlarged teeth on maxilla: present (0); absent (1).

**(87) Diastema between premaxillary and maxillary teeth: absent (0); present (1). (Liu et al., 2013: 26)**

(88) Maxillary tooth row: restricted to a level in front of the posterior margin of orbit (0); extending backward to a level below the posterior corner of orbit and/or the anterior corner of upper temporal fossa (1); extending backward to a level below the anterior one third to one half of upper temporal fossa (2).

**(89) Vomerine teeth: present (0); absent (1). (Liu et al., 2013: 28)**

**(90) Palatine teeth: four or more (0); one to three (1); absent (2). (Wang et al., 36)**

(91) Teeth on pterygoid flange: present (0); absent (1).

(92) Vertebrae: notochordal (0); non-notochordal (1).

(93) Vertebrae: amphicoelous (0); platycoelous (1); or other (2).

(94) Vertebral centrum: distinctly constricted in ventral view (0); with parallel lateral edges (1).

(95) Subcentral foramina: absent (0); present (1).

(96) Zygosphene-zygantrum articulation: absent (0); present (1).

(97) Zygapophyseal pachyostosis: absent (0); present (1).

**(98) Number of cervical vertebrae: 10 to 30 (0); more than 30 (1); fewer than 10 (2).**

(99) Cervical centra: rounded ventrally (0); keeled ventrally (1).

(100) Parapophysis: not shifting backward on centrum along cervical vertebral column (0); shifting backward on centrum along cervical vertebral column (1).

(101) Cervical intercentra: present (0); absent (1).

**(102) Number of dorsal vertebrae: 15 or fewer (0); 16 to 20 (1); 21 to 30 (2); 31 or more (3). (Wang et al., 2019: 74; Rieppel and Lin, 1995: 30)**

**(103) Hyposphere-hypantrum articulation absent (0); or present (1). (Wang et al., 2019: 75)**

**(104) Dorsal vertebrae neural spine height: less than double neural spine anteroposterior length (0); at least double neural spine anteroposterior length (1). (Liu et al., 2013: 31)**

(105) Distal articular surface on transverse processes of dorsal vertebrae: oblong (0); evenly rounded (1).

(106) Transverse processes of neural arches in dorsal region: relatively short (0); distinctly elongated (1).

(107) Distal end of transverse processes of dorsal vertebrae: not increasing in diameter (0); distinctly thickened (1).

(108) Sutural facets receiving pedicels of neural arch on dorsal surface of centrum in dorsal region: narrow (0); expanded into a cruciform or ‘butterfly-shaped’ platform (1).

(109) Dorsal intercentra: present (0); absent (1).

(110) Anteroposterior trend of increasing inclination of pre- and postzygapophyses within dorsal and sacral region: absent (0); present (1).

(111) A distinct free anterior process of cervical ribs: absent (0); present (1).

(112) Pachyostosis of dorsal ribs: absent (0); present (1);

**(113) Uncinate process on rib: absent (0); present (1). (New character)**

**(114) The last dorsal rib: longer (0); shorter (1) than the first sacral rib. (New character)**

(115) Number of sacral ribs: two (0); three (1); four or more (2).

(116) Distinct expansion of distal head of sacral ribs: present (0); absent (1).

(117) Sacral (and caudal) ribs or transverse processes and their respective centrum: sutured (0); fused (1).

**(118) Neural spine anticlination in tail: absent (0); present (1). (Ji et al., 2016: 153)**

**(119) Caudal ribs of proximal tail region: shorter or similar to sacral ribs (0); obviously longer than sacral ribs (1). (New character)**

**(120) Chevron: simple morphology, y-shaped (0); complex morphology with anteroposterior expansion on distal end. (Wang et al., 2019: 63)**

(121) Mineralized sternum: absent (0); present (1).

**(122) Gastralia segments: paired (0); three (1) or five (2). (Rieppel and Lin, 1995: 34)**

(123) Median gastral element: angulated (0); straight (1).

(124) The medial gastral rib element: with a single lateral process (0); with two-pronged lateral process (1).

**(125) Lateral gastral ribs without (0), or with (1) distinct angulation. (Wang et al., 2019: 80)**

(126) Cleithrum: present (0); absent (1).

(127) Clavicles: broad medially and generally slender (0); narrow medially and generally broad (1).

(128) Clavicles: not meeting in front of interclavicle (0); meeting in an interdigitating anteromedial suture (1).

(129) Anterolaterally expanded corners of clavicles: absent (0); present without distinct anterolateral process (1); present with distinct anterolateral process (2).

(130) Clavicle: applied to anterior (lateral) surface of scapula (0); applied to medial surface of scapula (1).

(131) Relationship between clavicles and interclavicle: in simple overlapping contact (0); anteromedioventral end of clavicle embracing lateral tip of interclavicle in a complex contact (1).

(132) Interclavicle: rhomboidal (0); T-shaped (1).

(133) Posterior process on (T-shaped) interclavicle: elongate (0); short (1); rudimentary or absent (2).

(134) Scapula: represented by a broad blade of bone (0); with a constriction separating a ventral glenoidal portion from a posteriorly directed dorsal wing (1); rod-like (2).

(135) Dorsal wing or process of eosauropterygian scapula: tapers to a blunt tip (0); ventrally expanded at its posterior end (1).

(136) Supraglenoid buttress: present (0); absent (1).

(137) Number of coracoid ossifications: one (0); two (1).

(138) Coracoid: of rounded contours (0); slightly waisted (1); strongly waisted (2); with expanded medial symphysis and ridge-like thickening of the bone extending from glenoid facet posteriorly along lateral edge of the bone, coracoid foramen not enlarged (3); with expanded medial symphysis and ridge-like thickening of the bone extending from glenoid facet transversely through the bone, coracoid foramen much enlarged (4).

(139) Coracoid foramen: enclosed by coracoid ossification (0); between coracoid and scapula (1).

(140) Pectoral fenestration: absent (0); present (1).

(141) Limbs: short and stout (0); long and slender (1).

(142) Foot: short and broad (0); long and slender (1).

(143) Humerus: rather straight (0); ‘curved’ (1).

**(144) Humerus length: shorter than or equal to femur (0); longer than femur (1). (New character)**

**(145) Humerus without (0) or with (1) vermiculate surface. (Rieppel and Lin, 1995: 40)**

(146) Deltopectoral crest: well-developed (0); reduced (1); absent (2).

(147) Insertional crest for latissimus dorsi muscle: prominent (0); reduced (1).

(148) Epicondyles of humerus: prominent (0); reduced (1).

(149) Ectepicondylar groove: open and notched anteriorly (0); open without anterior notch (1); closed (2); absent (3).

(150) Entepicondylar foramen: present (0); absent (1).

(151) Radius: shorter than ulna (0); longer than ulna (1); approximately of same length (2).

(152) Distal end of ulna: not expanded (0); distinctly expanded to at least the width of the proximal part (1).

(153) Total number of carpal ossifications: more than three (0); three (1); two (2).

**(154) Intermedium carpal: rounded (0) rectangular elongated at the preaxial margin of the distal tip of the ulna (1). (Rieppel and Lin, 1995: 44)**

(155) Iliac blade: well-developed (0); reduced but projecting beyond level of posterior margin of acetabular portion of ilium (1); reduced and no longer projecting beyond posterior margin of acetabular portion of ilium (2); absent, i.e., reduced to simple dorsal stub (3); elongated shaft (4).

**(156) Pubis: generally round (0); with concave anterior margin (1). (New character)**

(157) Pubis: with convex ventral (medial) margin (0); with concave ventral (medial) margin (1).

**(158) Obturator foramen in adult: present and closed (0); present and open (1); absent (2). (Added a new state)**

**(159) Ischium: generally round (0); with concave posterior margin (1). (New character)**

**(160) Thyroid fenestra: absent (0); present (1) (Wang et al., 2019: 77).**

(161) Acetabulum: oval (0); circular (1).

(162) Femoral shaft: stout and straight (0); slender and sigmoidally curved (1).

(163) Internal trochanter: well-developed (0); reduced (1).

(164) Intertrochanteric fossa: deep (0); distinct but reduced (1); rudimentary or absent (2).

(165) Distal femoral condyles: prominent (0); not projecting markedly beyond shaft (1).

(166) Anterior femoral condyle relative to posterior condyle: larger and extending further distally (0); smaller/equisized and of subequal extent distally (1).

**(167) Fibula: slender (0); distinctly expanded and wider than tibia (1). (modified from Liu et al., 2013: 40)**

(168) Total number of tarsal ossifications: four or more (0); three (1); two or fewer (2).

(169) Perforating artery: passes between astragalus and calcaneum (0); between distal heads of tibia and fibula proximal to astragalus (1).

(170) Proximal concavity of astragalus: absent (0); present (1).

(171) Calcaneal tuber: absent (0); present (1).

(172) Distal tarsal 1: present (0); absent (1).

(173) Distal tarsal 5: present (0); absent (1).

(174) Metatarsal 5: long and slender (0); distinctly shorter than other metatarsals and with a broad base (1).

(175) Metatarsal 5: straight (0); ‘hooked’ (1).

**(176) Hyperphalangy absent (0) or present (1) in manus. (Rieppel and Lin, 1995: 46)**

**(177） Ungual phalange in pes: long and sharp (0); short and blunt but not expanded (1); blunt and expanded, wider than the articulated proximal phalange (2). (New character)**

(178) Dermal armour ("osteoderms"): absent (0); present (1); forming single carapace, excluding endoskeletal elements (2); same as (2) but forming distinctly separate dorsal and pelvic carapaces (3); forming carapace, including endoskeletal elements (4). Note: this character has been changed to include the fact that some placodont taxa have a separate pelvic carapace.

(179) Distinctly open L-shaped (boomerang-shaped) jugal: absent (0); present (1).

(180) Palatine dentition: multiple rows with small numerous teeth/denticles (0); single row with four or more teeth (1), single row with three to one teeth/tooth (2); absent (3).

**(181) Marginal teeth: with convex (0) or concave (1) lingual surface of crown. (Cheng et al., 2016)**

Liu, J., Zhao, L. J., Li, C., et al. Osteology of *Concavispina biseridens* (Reptilia, Thalattosauria) from the Xiaowa Formation (Carnian), Guanling, Guizhou, China. Journal of Paleontology,

2013, 87(2): 341-350.

Neenan, J. M., Li, C., Rieppel, O., et al. The cranial anatomy of Chinese placodonts and the phylogeny of Placodontia (Diapsida: Sauropterygia). Zoological Journal of the Linnean Society, 2015, 175(2): 415-428.

Rieppel, O., Lin, K. B. Pachypleurosaurs (Reptilia: Sauropterygia) from the Lower Muschelkalk, and a review of the Pachypleurosauroidea. Field Museum of Natural History, 1995.

Scheyer, T. M., Neenan, J. M., Bodogan, T., et al. A new, exceptionally preserved juvenile specimen of *Eusaurosphargis dalsassoi* (Diapsida) and implications for Mesozoic marine diapsid phylogeny. Scientific Reports, 2017, 7(1): 4406.

Wang, W., Li, C., Scheyer, T. M., et al. A new species of *Cyamodus* (Placodontia, Sauropterygia) from the early Late Triassic of south-west China. Journal of Systematic Palaeontology, 2019, 17: 1457-1476.

#### 3. Specimen list of the sampled Triassic marine diapsids

##### Supplementary Table 1 Specimen list of sampled Triassic marine diapsids

| Taxa | Specimen number<br>(H: holotype)<br>(*main specimen for coding) | Observed by WW<br>(√) or cited from<br>literature |
|---|---|---|
| *Hanosaurus hupehensis* | IVPP V3231 (H)* | √ |
| | IVPP V15911* | √ |
| *Helveticosaurus zollingeri* | PIMUZ T4352 (H)* | √ |
| | PIMUZ T4353 | √ |
| | PIMUZ T4354 | √ |
| *Eusaurosphargis dalsassoi* | BES SC 390 (H) | √ |
| | PIMUZ A/III 4380* | √ |
| *Sinosaurosphargis yunguiensis* | IVPP V 17040 (H)* | √ |
| | IVPP V 16076 | √ |
| | ZMNH M 8797 | √ |
| *Largocephalosaurus polycarpon* | WIGM SPC V 1009 (H) | Cheng et al., 201 |
| *Largocephalosaurus qianensis* | IVPP V 15638 (H)* | √ |
| | GMPKU-P-1532-A | √ |
| | GMPKU-P-1532-B | √ |
| *Atopodentatus unicus* | WIGM SPC V 1107 (H)* | Cheng et al., 2014 |
| | IVPP V 20191* | √ |
| | IVPP V 20292 | √ |
| *Palatodonta bleekeri* | TW480000470 (H)* | Neenan et al., 2013 |
| *Paraplacodus broilii* | PIMUZ T2806 (H)* | √ |
| | PIMUZ T4773 | √ |
| | PIMUZ T4776 | √ |
| | PIMUZ T4827* | √ |
| | PIMUZ T5847 | √ |
| | BSP 1953 XV 5 | √ |
| *Pararcus diepenbroeki* | TWE 480000454 (H)* | Klein and Scheyer, 2013 |
| *Placodus gigas* | BSP AS VII 1208 (H)* | √ |
| | UMO BT13 | Sues, 1987 |
| | BSP 1968 I 75 | √ |
| | SMF R-I035* | √ |
| *Placodus inexpectatus* | GMPKU-P-1054 | √ |
| | IVPP V 14996 | √ |
| *Cyamodus rostratus* | UMO BT 748 (H) | Rieppel, 2001 |

| | | |
|---|---|---|
| | SMNS 17403 | √ |
| | UMO BT 2172 | Rieppel, 2001 |
| *Cyamodus kuhnschnyderi* | SMNS 15855 (H) | √ |
| | SMNS 16270 | √ |
| | SMNS 18380 | √ |
| *Cyamodus muensteri* | BSP AS VII 1210 (H) | √ |
| *Cyamodus hildegardis* | PIMUZ T4763 (H) | √ |
| | PIMUZ T58 | √ |
| | PIMUZ T4768 | √ |
| | PIMUZ T4771 | √ |
| *Cyamodus orientalis* | ZMNH M8820 (H)* | √ |
| *Sinocyamodus xinpuensis* | IVPP V 11872 (H) | √ |
| | IVPP V17051* | √ |
| *Parahenodus atancensis* | MUPA ATZ0104 (H)* | de Miguel et al., 2018 |
| *Henodus chelyops* | GPIT Specimen II (H)* | √ |
| | GPIT Specimen I~VIII | √ |
| *Macroplacus raeticus* | BSP 1967 I 324 (H) | √ |
| *Protenodontosaurus italicus* | MFSN 1819GP | Rieppel, 2000 and 2001 |
| | MFSN 1923GP | |
| | SMNS 91423* | √ |
| *Psephochelys polyosteoderma* | IVPP V 12442* | √ |
| | EBGL009* | √ |
| *Placochelys placodonta* | FAFI Ob/2323/Vt.3 (H)* | Rieppel 2000 and 2001 |
| | MB.R.1765 | |
| *Psephoderma alpinum* | BSP AS I 8 (H) | √ |
| | MSNM V471* | √ |
| | MSNM V527* | √ |
| | PIMUZ A/III 1491 | √ |
| *Glyphoderma kangi* | ZMNH M 8729 (H)* | √ |
| *Anarosaurus heterodontus* | IGWH M4/12 (H) | Rieppel, 2000 |
| | NME 480000125* | Klein, 2009 and 2012 |
| | NME 480000127* | |
| | NME 480000130* | |
| *Dactylosaurus gracillis* | MGU Wr 3871s (H)* | Rieppel and Lin, 1995 |
| *Serpianosaurus mirigiolensis* | PIMUZ T3931 (H)* | √ |
| | PIMUZ T3086* | √ |
| | PIMUZ T1071* | √ |
| | PIMUZ T3689 | √ |
| | PIMUZ T3680 | √ |
| | PIMUZ T3042 | √ |
| | PIMUZ T3933 | √ |

| | | |
|---|---|---|
| *Neusticosaurus pusillus* | BMNH R53 (H) | Rieppel, 2000 |
| | PIMUZ T3934* | √ |
| | PIMUZ T3597* | √ |
| | PIMUZ T3661 | √ |
| | PIMUZ T3948 | √ |
| | PIMUZ T3576 | √ |
| *Neusticosaurus edwardsi* | PIMUZ T3437 | √ |
| | PIMUZ T3407 | √ |
| | PIMUZ T3439 | √ |
| | PIMUZ T3576 | √ |
| *Neusticosaurus peyer* | PIMUZ T3462 | √ |
| | PIMUZ T3705 | √ |
| | PIMUZ T3582 | √ |
| | PIMUZ T3403 | √ |
| *Odoiporosaurus teruzzii* | BES SC 1893 (H)* | √ |
| *Majiashanosaurus discocoracoidis* | AGM-AGB5954 (H)* | √ |
| *Wumengosaurus delicatomandibularis* | GMPKU-P-1210 (H) | Jiang et al., 2008 |
| | GMPKU-P-1209 | |
| | NMNS-KIKO-F071129-Z | Wu et al., 2011 |
| | IVPP V15314* | √ |
| | ZMNH M8758* | √ |
| *Qianxisaurus chajiangensis* | NMNS-KIKO-F044630 (H)* | Cheng et al., 2012 |
| *Diandongosaurus acutidentatus* | IVPP V17761 (H)* | √ |
| | NMNS-000933-F03498 | Sato et al., 2013 |
| | BGPDB-R0001 | Liu et al., 2015 |
| *Dianmeisaurus gracilis* | IVPP V18630 (H)* | √ |
| | IVPP V17054 | √ |
| *Dianopachysaurus dingi* | LPV 31365 (H)* | Liu et al., 2011 |
| *Dawazisaurus brevis* | NMNS000933-F034397 | Cheng et al., 2016 |
| *Keichousaurus hui* | IVPP V952 (H) | √ |
| | IVPP V17047* | √ |
| | IVPP V17043 | √ |
| | IVPP V16920 | √ |
| | NMNS-cyn-2003-25* | Holmes et al., 2008 |
| | NMNS-cyn-2005-05 | |
| | NMNS-cyn-2005-12 | |
| | NMNS-cyn-2005-15 | |
| | NMNS-cyn-2005-18 | |
| | NMNS-cyn-2005-24 | |
| *Panzhousaurus rotundirostris* | GMPKU-P-1059 (H) | Jiang et al., 2019 |
| *Simosaurus gaillardoti* | MNHN AC.9028 (H) | Rieppel, 2000 |
| | SMNS 16700 | √ |
| | SMNS 10360* | √ |

| | | |
|---|---|---|
| | SMNS 15860 | √ |
| | SMNS16767 | √ |
| | SMNS 50714* | √ |
| | SMNS 50715 | √ |
| *Paludidraco multidentatus* | MUPA-ATZ0101 (H)* | de Miguel, et al., 2018 |
| *Germanosaurus schafferi* | NHMW unnumbered (H)* | Rieppel, 2000 |
| *Nothosaurus youngi* | IVPP V 13590 (H)* | √ |
| | WS-30-R24* | Ji et al., 2014 |
| *Lariosaurus xingyiensis* | IVPP V11866 (H)* | √ |
| | XNGM-WS-30-R9* | Lin et al., 2017 |
| *Corosaurus alcovensis* | UW 5484 (H)* | Storrs, 1991; Rieppel, 2000 |
| | FMNH PR480* | |
| | FMNH PR1369 | |
| | YPM 41031 | |
| *Cymatosaurus fridericianus* | IGWH unnumbered (H) | Rieppel, 2000 |
| | BGR S 44/3 | |
| *Augustasaurus hagdorni* | FMNH PR 1774 (H) | Sander et al., 1997 |
| *Pistosaurus longaevus* | UMO unnumbered (H)* | Rieppel, 2000 |
| | SMF R4041* | √ |
| *Yunguisaurus liae* | NMNS 004529/F003862 (H)* | Cheng et al., 2006 |
| | ZMNH M8738* | √ |
| | IVPP V14993 | √ |
| *Wangosaurus brevirostris* | GMPKU-P-1529 (H)* | √ |
| *Bobosaurus forojuliensis* | MFSN 27285 | Dalla Vecchia, 2006 |
| *Miodentosaurus brevis* | NMNS 004727/F003960 (H)* | Cheng et al., 2007 |
| | ZMNH M8742* | √ |
| *Anshunsaurus huangguoshuensis* | IVPP V11835 (H)* | √ |
| | IVPP V11834* | √ |
| | GMPKU 2000-028 | √ |
| *Xinpusaurus suni* | IVPP V11860* | √ |
| | IVPP V12673* | √ |
| | IVPP V14372 | √ |
| | Gmr 101 | Liu, 2013 |
| *Concavispina biseridens* | ZMNH M8804 (H)* | √ |
| *Nanchangosaurus suni* | GMC V646 (H) | Wang, 1959 |
| | SSTM 5025* | √ |
| | WGSC 26006* | Chen et al., 2014a |
| *Eohupehsuchus brevicollis* | WGSC V26003* | Chen et al., 2014b |
| *Hupehsuchus nanchangensis* | IVPP V3232 (H)* | √ |
| | IVPP V 4068 | √ |
| | WGSC 26004 | Chen et al., 2014c |

| | | |
|---|---|---|
| | ZMNH M8217* | √ |
| *Parahupehsuchus longus* | WGSC 26005 (H) | Chen et al., 2014c |
| *Eretmorhipis carrolldongi* | WGSC 26020 (H)* | Chen et al., 2015 |
| | IVPP V4070* | √ |
| *Cartorhynchus lenticarpus* | AGB6257 (H)* | √ |
| *Sclerocormus parviceps* | AGB6265 (H)* | √ |
| *Chaohusaurus geishanensis* | IVPP V 4001 | √ |
| | IVPP V 11361 | √ |
| | IVPP V V11362 | √ |
| | AGM P45-H85-25* | √ |
| | AGM P45-H85-20* | √ |
| | AGM P45-H85-24 | √ |
| | AGM-MT10010 | √ |
| *Pappochelys rosinae* | SMNS 91360 (H)* | √ |
| | SMNS 90013* | √ |
| *Eorhynchochelys sinensis* | SMMP 000016 (H)* | √ |
| *Odontochelys semitestacea* | IVPP V15639 (H)* | √ |
| | IVPP V13240* | √ |
| | IVPP V15653 | √ |

Cheng, Y. N., Wu, X. C., Sato, T., et al. A new thalattosaurian (Reptilian: Diapsida) from the Upper

Triassic of Guizhou, China. Vertebrata Palasiatica, 2007, 45: 246-260.

Cheng, Y. N., Wu, X. C., et al. A new eosauropterygian (Diapsida, Sauropterygia) from the Triassic of China. Journal of Vertebrate Paleontology, 2012, 32(6): 1335-1349.

Cheng, Y. N., Wu, X. C., Sato, T., et al. *Dawazisaurus brevis*, a new eosauropterygian from the middle Triassic of Yunnan, China. Acta Geologica Sinica (English Edition): 2016, 90(2), 401-424.

Dalla, Vecchia., F. M.. A new sauropterygian reptile with plesiosaurian affinity from the Late Triassic of Italy. Rivista Italiana di Paleontologia e Stratigrafia (Research In Paleontology and Stratigraphy): 2006,112(2).

de Miguel Chaves, C., Ortega, F., Pérez-García, A. A new placodont from the Upper Triassic of Spain provides new insights on the acquisition of the specialized skull of Henodontidae. Papers in Palaeontology, 2018, 4(4): 567-576.

de Miguel Chaves, C., Ortega, F., Pérez-García, A. New highly pachyostotic nothosauroid interpreted as a filter-feeding Triassic marine reptile. Biology Letters, 2018, 14(8): 20180130.

Holmes, R., Cheng, Y. N., Wu, X. C.. New information on the skull of *Keichousaurus hui* (Reptilia: Sauropterygia) with comments on sauropterygian interrelationships. Journal of Vertebrate Paleontology, 2008, 28(1): 76-84.

Ji, C., Jiang, D. Y., Rieppel, O., et al. A new specimen of *Nothosaurus youngi* from the Middle Triassic of Guizhou, China. Journal of Vertebrate Paleontology, 2014, 34(2): 465-470.

Jiang, D. Y., O. Rieppel, R. Motani, et al.. A new middle Triassic eosauropterygian (Reptilia, Sauropterygia) from southwestern China. Journal of Vertebrate Paleontology, 2008, 28:1055–1062.

Jiang, D. Y., Lin, W. B., Rieppel, O., et al. A new Anisian (Middle Triassic) eosauropterygian (Reptilia, Sauropterygia) from Panzhou, Guizhou Province, China[J]. Journal of Vertebrate Paleontology, 2019, 38(4): (1)-(9).

Klein, N. Skull morphology of *Anarosaurus heterodontus* (Reptilia: Sauropterygia: Pachypleurosauria) from the Lower Muschelkalk of the Germanic Basin (Winterswijk, The Netherlands). Journal of Vertebrate Paleontology, 2009, 29(3): 665-676.

Klein, N. Postcranial morphology and growth of the pachypleurosaur *Anarosaurus heterodontus* (Sauropterygia) from the Lower Muschelkalk of Winterswijk, The Netherlands. Paläontologische Zeitschrift, 2012, 86(4): 389-408.

Klein, N, Scheyer, T. M. A new placodont sauropterygian from the Middle Triassic of the Netherlands. Acta Palaeontologica Polonica, 2013, 59(4): 887-903.

Lin, W. B., Jiang, D. Y., Rieppel, O., et al. A New Specimen of *Lariosaurus xingyiensis* (Reptilia, Sauropterygia) from the Ladinian (Middle Triassic) Zhuganpo Member, Falang Formation, Guizhou, China. Journal of Vertebrate Paleontology, 2017, 37(2): e1278703.

Liu, J., Rieppel, O., Jiang, D. Y., et al. A new pachypleurosaur (Reptilia: Sauropterygia) from the lower Middle Triassic of southwestern China and the phylogenetic relationships of Chinese pachypleurosaurs. Journal of Vertebrate Paleontology, 2011, 31(2): 292-302.

Liu, X. Q., Lin, W. B., Rieppel, O. et al. A new specimen of *Diandongosaurus acutidentatus* (Sauropterygia) from the Middle Triassic of Yunnan, China. Vertebr Palasiatica:2015, 10, 281-290.

Neenan, J. M., Klein, N., Scheyer, T. M. European origin for sauropterygian marine reptiles and the evolution of placodont crushing dentition. Nature Communications, 2013, 4: 1621.

Sues, H. D. On the skull of *Placodus gigas* and the relationships of the Placodontia. Journal of Vertebrate Paleontology, 1987, 7(2): 138-144.

Rieppel, O. Cranial anatomy of *Placochelys placodonta* Jaekel, 1902, and a review of the Cyamodontoidea (Reptilia, Placodonta). Fieldiana (Geology) n.s., 2001 45: 1-104.

Rieppel, O. Sauropterygia I, Placodontia, Pachypleurosauria, Nothosauroidea, Pistosauroidea. In: Wellnhofer P, ed. Encyclopedia of paleoherpetology. Munich: Verlag Dr. Friedrich Pfeil, 2000: 1-134.

Rieppel, O., Lin, K. B. *Pachypleurosaurs* (Reptilia: Sauropterygia) from the Lower Muschelkalk, and a review of the Pachypleurosauroidea (No. 32). Field Museum of Natural History, 1995.

Sato, T., Cheng, Y. N., Wu, X. C., et al. *Diandongosaurus acutidentatus* Shang, Wu & Li, 2011 (Diapsida: Sauropterygia) and the relationships of Chinese eosauropterygians. Geological Magazine, 2014, 151(1): 121-133.

Sander, P. M., Rieppel, O., Bucher, H. A new pistosaurid (Reptilia: Sauropterygia) from the Middle Triassic of Nevada and its implications for the origin of the plesiosaurs. Journal of Vertebrate Paleontology, 1997, 17(3): 526-533.

Storrs, G. W. Anatomy and Relationships of *Corosaurus alcovensis* (Diapsida: Sauropterygia) and the Triassic Alcova Limestone of Wyoming. Bulletin of the Peabody Museum of Natural History, 1991, 44:1-151.

Wang, G. M. A new marine reptile from Hupeh Province. ActaPalaeontologica Sinica, 1959, 7 (5): 367-373. (in Chinese)

#### 4. Phylogenetic results and clade synapomorphies

The monophyly of Sauropterygiformes, including saurosphargids, placodontiforms, eosauropterygians, and relevant reptiles, is confirmed, and its internal relationships are well resolved (Fig. 2; Supplementary Fig. 5-10). Saurosphargidae, a group composed of *Eusaurosphargis*, *Largocephalosaurus*, and *Sinosaurosphargis*, is confirmed to be the sister group of Sauropterygia, despite the interpositions of the enigmatic and unstable taxa *Helveticosaurus* and *Atopodentatus*. Sauropterygia bifurcates into Placodontiformes and Eosauropterygia. Placodonts are indubitable sauropterygians, but not diverged as early as previously considered within sauropterygians and are gradually specialized through time after some conical-toothed and unarmored pioneers. Placodontiformes is maintained and its intra-relationships are well resolved including the monophyletic Cyamodontoidea and more derived Placochelyida developing heavy armors and carapaces. Our topology of placodontiforms is well in accordance with other studies for placodonts, though there are a few minor differences such as here placing the highly specialized henodontids as the most derived taxa. Notably, the monophyletic 'Pachypleurosauria' is not conclusively recovered here like most recent analyses, despite incomplete sampling of non-eusauropterygian eosauropterygians in these previous studies. In Eosauropterygia, *Wumengosaurus* and *Qianxisaurus* are the basal-most members, whereas the other Chinese pachypleurosaur-like taxa are more derived within eosauropterygians showing close affinities with each other, but whether these Chinese pachypleurosaur-like taxa form a monophyletic group is unsolved. European pachypleurosaurs form a monophyletic lineage as the family Pachypleurosauridae following the previous definition from Rieppel. Eusauropterygia is recovered including all the large eosauropterygians (adult length > 1 m), in which Simosauridae, Nothosauridae, and Pistosauroidea are monophyletic as previously defined in the studies of Rieppel, Neenan, and other colleagues.

Based on the strict consensus results (Fig. 2) involving 54 taxa and 181 characters (Supplementary Fig. 8), the definitions and synapomorphies of crucial clades are listed as following (Char. is the abbreviation of Character):

**Thalattosauria** Merriam, 1904

Definition: the most recent common ancestor of *Askeptosaurus* and *Xinpusaurus* and all of its descendants.

Synapomorphies: nasal reduced (Char. 11: 0→1); postfrontal and postorbital fused (Char. 28: 0→1); upper temporal fossa secondarily closed (Char. 42: 0→4); pubis with concave anterior margin (Char. 156: 0→1); fibula distinctly expanded and wider than tibia (Char. 167: 0→1); proximal concavity of astragalus present (Char. 170: 0→1).

**Unnamed clade** (Ichthyosauriformes + Sauropterygiformes)

Synapomorphies: distinct coronoid process of lower jaw absent (Char. 72: 1→0); coracoid foramen between coracoid and scapula (Char. 139: 0→1); obturator foramen in adult present and open (Char. 158: 0→1); distinctly open L-shaped jugal present (Char. 179: 0→1).

**Ichthyosauriformes** Motani et al., 2015

Definition: the most recent common ancestor of *Cartorhynchus* and *Ichthyosaurus* and all of its descendants.

Synapomorphies: nasal longer than frontal (Char. 10: 0→1); nasal anteriorly extending beyond external naris (Char. 12: 0→1); parietal skull table weakly constricted (Char. 34: 0→1); durophagous dentition present (Char. 82: 0→1); vertebral centrum with parallel lateral edges (Char. 94: 0→1); tarsal ossifications are two or less (Char. 168: 0→2).

**Sauropterygiformes** Wang et al., 2021

Definition: the most recent common ancestor of *Hanosaurus* and Plesiosauria and all of its descendants.

Synapomorphies: palate akinetic (Char. 62: 0→1); gastralia segments as five (Char. 122: 0/1→2); clavicles narrow medially (Char. 127: 0→1); pectoral fenestration present (Char. 140: 0→1); iliac blade reduced but projecting beyond level of posterior margin of acetabular portion of ilium (Char. 155: 0→1); marginal teeth with concave lingual surface of crown (Char. 181: 0→1).

**Saurosphargidae** Li et al., 2011

Definition: the most recent common ancestor of *Eusaurosphargis*, *Largocephalosaurus*, and *Sinosaurosphargis* and all of its descendants.

Synapomorphies: splenial bone excluded from the mandibular symphysis (Char. 71: 0→1); uncinate process on ribs present (Char. 113: 0→1); dermal armor present (Char. 178: 0→1).

**Sauropterygia** Owen, 1860

Definition: the most recent common ancestor of Placodontiformes and Eosauropterygia and all of its descendants.

Synapomorphies: humerus curved (Char. 143: 0→1); femoral shaft slender and sigmoidally curved (Char. 162: 0→1); marginal teeth with convex lingual surface of crown (Char. 181: 1→0).

**Placodontiformes** Neenan et al., 2013

Definition: the most recent common ancestor of *Palatondonta* and *Henodus* and all of its descendants.

Synapomorphies: squamosal broadly separated from ventral margin of skull (Char. 45: 0→1); pterygoid shorter than palatine (Char. 66: 0→1); mandibular symphysis somewhat enforced (Char. 74: 0→1); anterior (premaxillary and dentary) teeth strongly procumbent (Char. 84: 0→1).

**Eosauropterygia** Rieppel, 1994

Definition: the most recent common ancestor of *Wumengosaurus* and Plesiosauria and all of its descendants.

Synapomorphies: preorbital region distinctly longer than postorbital region (Char. 2: 0→1); transverse process of neural arches in dorsal region relatively short (Char. 106: 1→0); coracoid strongly waisted (Char. 138: 0→2/3); entepidcondylar foramen present (Char. 150: 1→0).

**Pachypleurosauridae** Nopcsa, 1928

Definition: the most recent common ancestor of *Dactylosaurus* and *Neusticosaurus* and all of its descendants.

Synapomorphies: dermatocranial bones relatively smooth (Char. 1: 0→1); postfrontal with reduced lateral process and hence more of an elongate shape (Char. 27: 1→2); dorsal wing of epipterygoid narrower than its base (Char. 54: 0→1); ectopterygoid absent (Char. 68: 0→1); distinct expansion of distal head of sacral ribs absent (Char. 116: 0→1); gastralia segment as three (Char. 122: 2→1); epicondyles of humerus prominent (Char. 148: 1→0);

**Eusauropterygia** Tschanz, 1989

Definition: the most recent common ancestor of *Simosaurus* and Plesiosauria and all of its descendants.

Synapomorphies: postorbital region distinctly larger than preorbital region (Char. 2: 0→2); upper temporal fossa present and distinctly larger than orbit (Char. 42: 3→2); splenial bone excluded from the mandibular symphysis (Char. 71: 0→1); trough on the dorsal surface of retroarticular process absent (Char. 78: 1→0); anterior (premaxillary and dentary) teeth strongly procumbent (Char. 84: 0→1); distinctly open L-shaped (boomerang-shaped) jugal absent (Char. 179: 1→0).

**Nothosauridae** Baur, 1889

Definition: The most recent common ancestor of *Nothosaurus* and *Lariosaurus* and all of its descendants.

Synapomorphies: nasals meeting in dorsomedial suture (Char. 13: 2→0); anterolateral processes of frontals reduced (Char. 22: 0→1); postfrontal with reduced lateral process and hence more of an elongated shape (Char. 27: 1→2); pineal foramen located within a deep trough (Char. 36: 0→1).

**Pistosauroidea** Baur, 1890

Definition: the most recent common ancestor of *Corosaurus* and Plesiosauria and all of its descendants.

Synapomorphies: occipital crest present but squamosals not meeting behind parietal (Char. 57: 2→1); distal end of transverse process of dorsal vertebrae distinctly thickened (Char. 107: 0→1); obturator foramen in adult absent (Char. 158: 1→2)

### 5. Taxon ages of Triassic marine reptiles

**Supplementary Table 2 Ages of Triassic marine reptiles**

| Taxon | First-appeared date (Ma) | Last-appeared date (Ma) |
|---|---|---|
| *Youngina* | 251.9 | 251.9 |
| *Cartorhynchus* | 249.9 | 246.7 |
| *Sclerocormus* | 249.9 | 246.7 |
| *Chaohusaurus* | 249.9 | 246.7 |
| *Miodentosaurus* | 237.0 | 227.3 |
| *Anshunsaurus* | 241.5 | 227.3 |
| *Xinpusaurus* | 241.5 | 227.3 |
| *Concavispina* | 237.0 | 227.3 |
| *Helveticosaurus* | 241.5 | 241.5 |
| *Eusaurosphargis* | 241.5 | 241.5 |
| *Sinosaurosphargis* | 246.7 | 241.5 |
| *Largocephalosaurus* | 246.7 | 241.5 |
| *Atopodentatus* | 246.7 | 241.5 |
| *Palatodonta* | 246.7 | 241.5 |
| *Paraplacodus* | 241.5 | 241.5 |
| *Pararcus* | 246.7 | 241.5 |
| *Placodus* | 246.7 | 237.0 |
| *Cyamodus* | 246.7 | 227.3 |
| *Sinocyamodus* | 237.0 | 227.3 |
| *Parahenodus* | 237.0 | 209.5 |
| *Henodus* | 237.0 | 227.3 |
| *Macroplacus* | 209.5 | 201.4 |
| *Protenodontosaurus* | 237.0 | 227.3 |
| *Psephochelys* | 237.0 | 227.3 |
| *Placochelys* | 237.0 | 201.4 |
| *Psephoderma* | 227.3 | 201.4 |
| *Glyphoderma* | 241.5 | 237.0 |
| *Anarosaurus* | 246.7 | 241.5 |
| *Dactylosaurus* | 246.7 | 241.5 |
| *Serpianosaurus* | 241.5 | 241.5 |
| *Neusticosaurus* | 241.5 | 237.0 |
| *Odoiporosaurus* | 246.7 | 241.5 |
| *Hanosaurus* | 249.9 | 246.7 |
| *Majiashanosaurus* | 249.9 | 246.7 |
| *Wumengosaurus* | 246.7 | 241.5 |

| | | |
|---|---|---|
| *Qianxisaurus* | 241.5 | 237.0 |
| *Diandongosaurus* | 246.7 | 241.5 |
| *Dianmeisaurus* | 246.7 | 241.5 |
| *Dianopachysaurus* | 246.7 | 241.5 |
| *Dawazisaurus* | 246.7 | 241.5 |
| *Keichousaurus* | 241.5 | 237.0 |
| *Panzhousaurus* | 246.7 | 241.5 |
| *Simosaurus* | 241.5 | 237.0 |
| *Paludidraco* | 227.3 | 227.3 |
| *Germanosaurus* | 246.7 | 241.5 |
| *Nothosaurus* | 246.7 | 237.0 |
| *Lariosaurus* | 246.7 | 237.0 |
| *Corosaurus* | 249.9 | 241.5 |
| *Cymatosaurus* | 246.7 | 241.5 |
| *Augustasaurus* | 246.7 | 241.5 |
| *Pistosaurus* | 246.7 | 241.5 |
| *Yunguisaurus* | 241.5 | 237.0 |
| *Wangosaurus* | 241.5 | 237.0 |
| *Bobosaurus* | 237.0 | 227.3 |

### 6. Measurements of secondarily aquatic reptiles

Measurements are documented in the EXCEL file (reference listed blow).

Abbreviations and explanations of the Excel table: SL, skull (maximum) length; SW, skull (maximum) width; NL, neck length; BL, body or trunk length (including dorsal vertebrae and sacrum); BW, body or trunk (maximum) width; TL, tail length; TW, tail (proximal/maximum) width; FL, forelimb length; HL, hind limb length. Data with '~' means the estimation due to the incompleteness or the preservation; specimen number with '*' means that the measurement was partially based on the reconstructions. The data with no reference were collected from the specimen or photos taken by the authors. Alternatively, the data were collected from the literature or measured by us from the figures. The body width was estimated by doubling the distance between the trunk lateral margin (being bordered by ribs) and the body axis. The tail width was estimated by doubling the proximal caudal rib length plus the centrum width.

Supplementary Figures

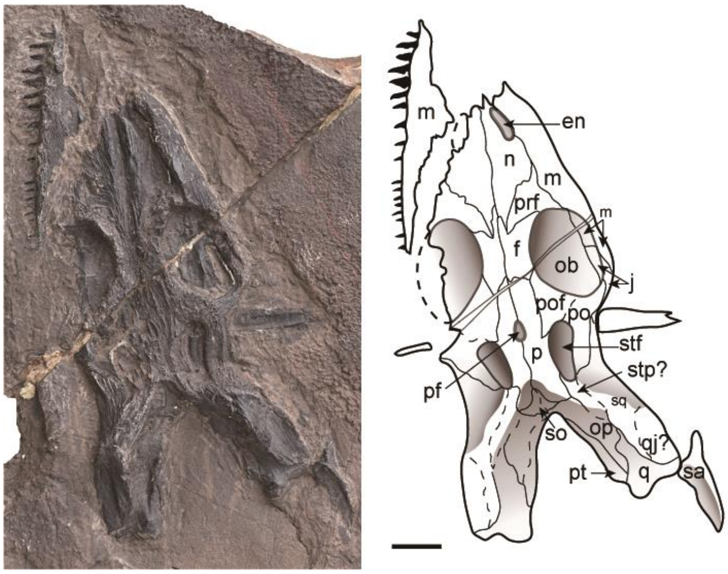


**Supplementary Fig. 1. Skull of the holotype of *Hanosaurus hupehensis* (IVPP V 3231).**

Abbreviations: en, external naris; f, frontal; m, maxilla; n, nasal; ob, orbit; op, opisthotic; p, parietal; pf, parietal foramen; po, postorbital; pof, postfrontal; prf, prefrontal; pt, pterygoid; q, quadrate; qj, quadratojugal; sa, surangular; so, supraoccipital; sq, squamosal; stf, supratemporal fenestra; stp, supratemporal. (scale bar=1 cm)

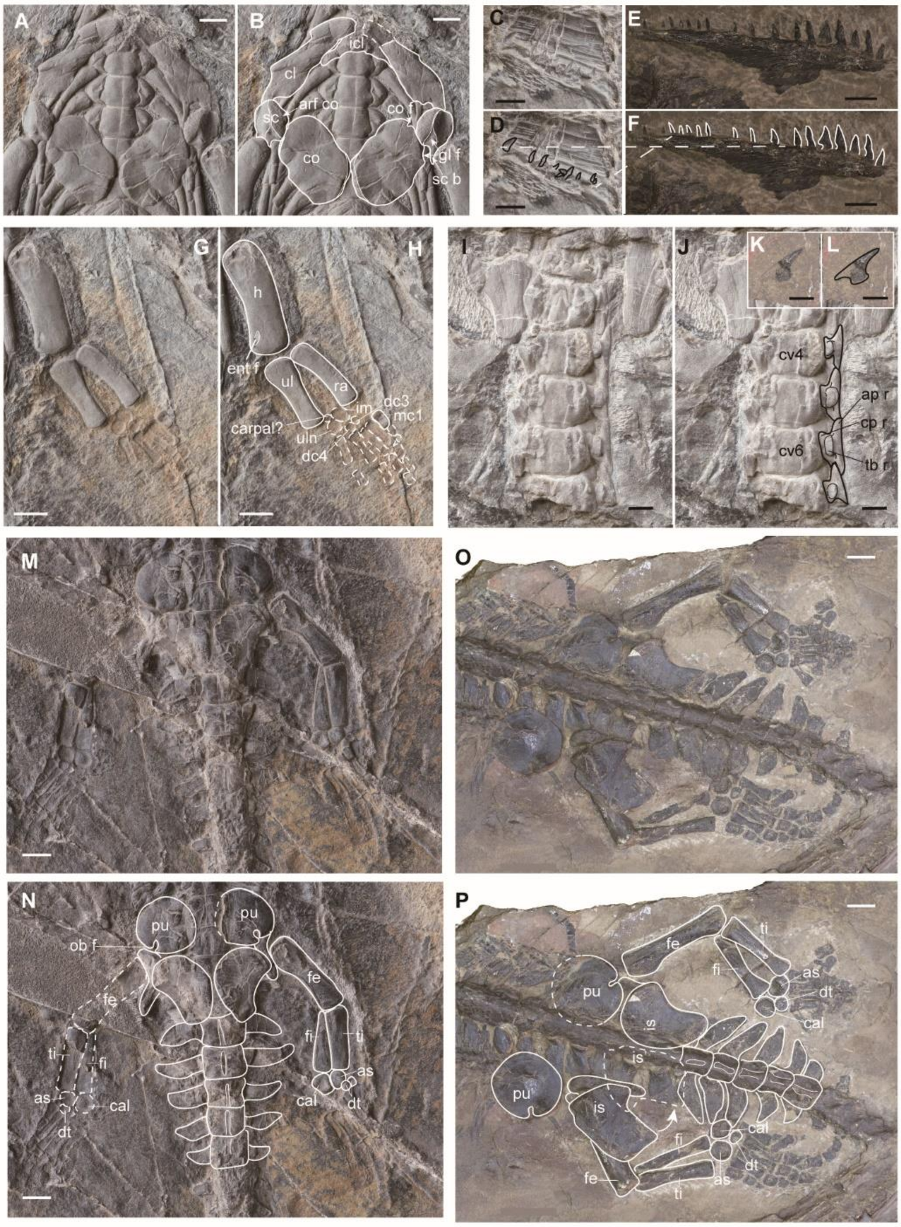


**Supplementary Fig. 2. Detailed comparisons between the new specimen (IVPP V 15911) and the holotype of *Hanosaurus hupehensis* (IVPP V 3231).**

(a-b), pectoral girdle of V 15911 (scale bar=1 cm); (c-d), dentition of V 15911 and (e-f), V 3231 (scale bar =5 mm); (g-h), left forelimb of V 15911 (scale bar=1 cm); (I-J), cervical rib(s) of V 15911 and (k-l), V 3231 (scale bar=5 mm); (m-n), pelvic girdle and hind limbs of V 15911 and (o-p), V 3231 (scale bar=2 cm). Abbreviations: ap r, anterior process of rib; arf co, articular surface on scapula to coracoid; as, astragalus; cal, calcaneum; co f, coracoid foramen; cp r, capitulum of rib; cv, cervical vertebra; dc, distal carpal; ent f, entepicondylar foramen; fe, femur; fi, fibula; gl f, glenoid fossa; h, humerus; im, intermedium; is, ischium; mc, metacarpal; ob f, obturator foramen; pu, pubis; ra, radius; sc b, scapula blade; tb r, tuberculum of rib; ti, tibia; ul, ulna; uln, ulnare.

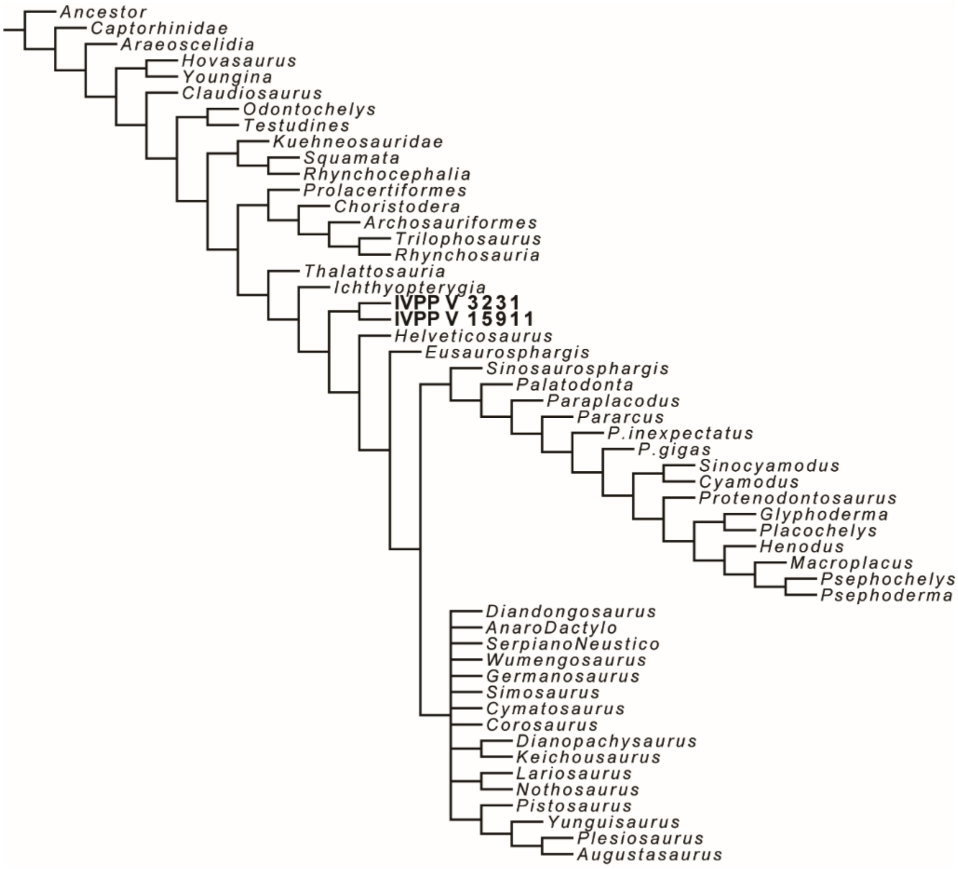


**Supplementary Fig. 3. Strict consensus tree based on the data matrix of Neenan et al. showing IVPP V 3231 and IVPP V 15911 as a single clade.**

618 steps, 5 most parsimony trees (MPTs), CI: 0.314, RI: 0.694.

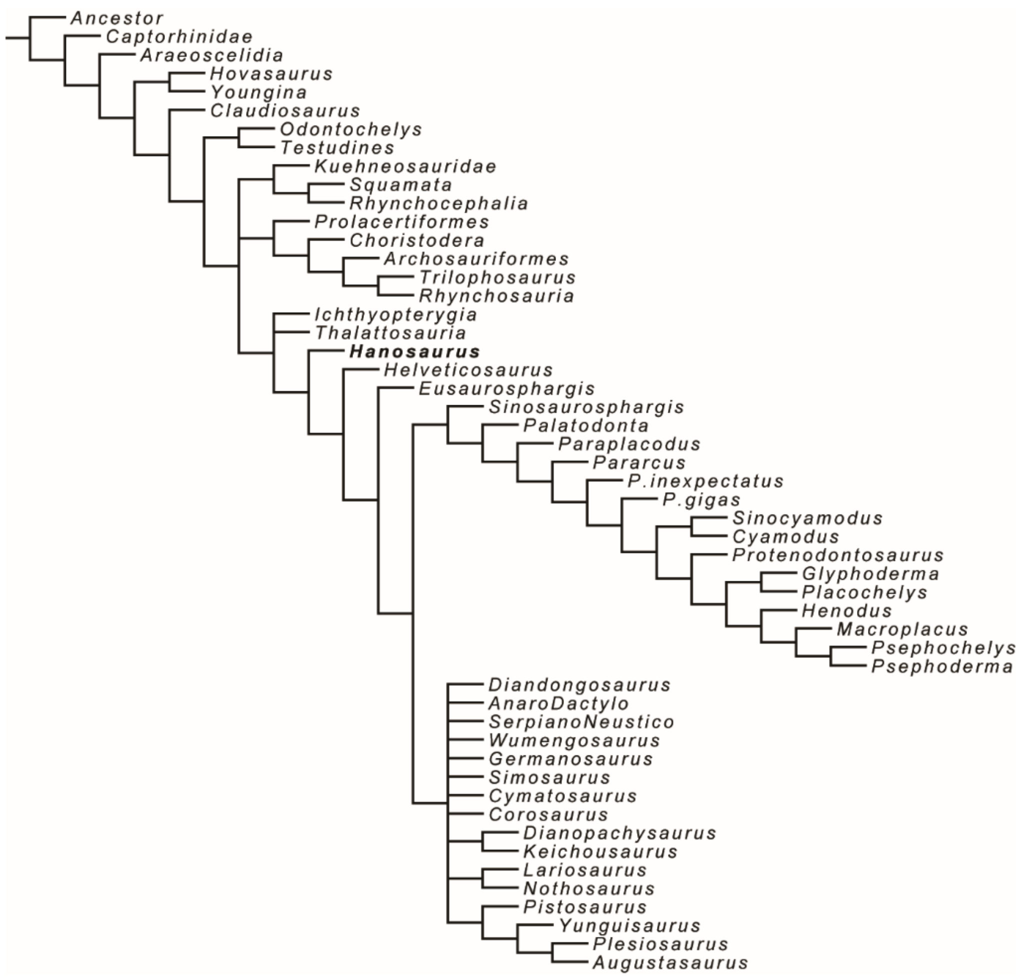


**Supplementary Fig. 4. Strict consensus tree based on the data matrix of Neenan et al. showing *Hanosaurus* as the basal-most taxon of sauropterygiforms.**

615 steps, 15 MPTs, CI: 0.315, RI: 0.695.

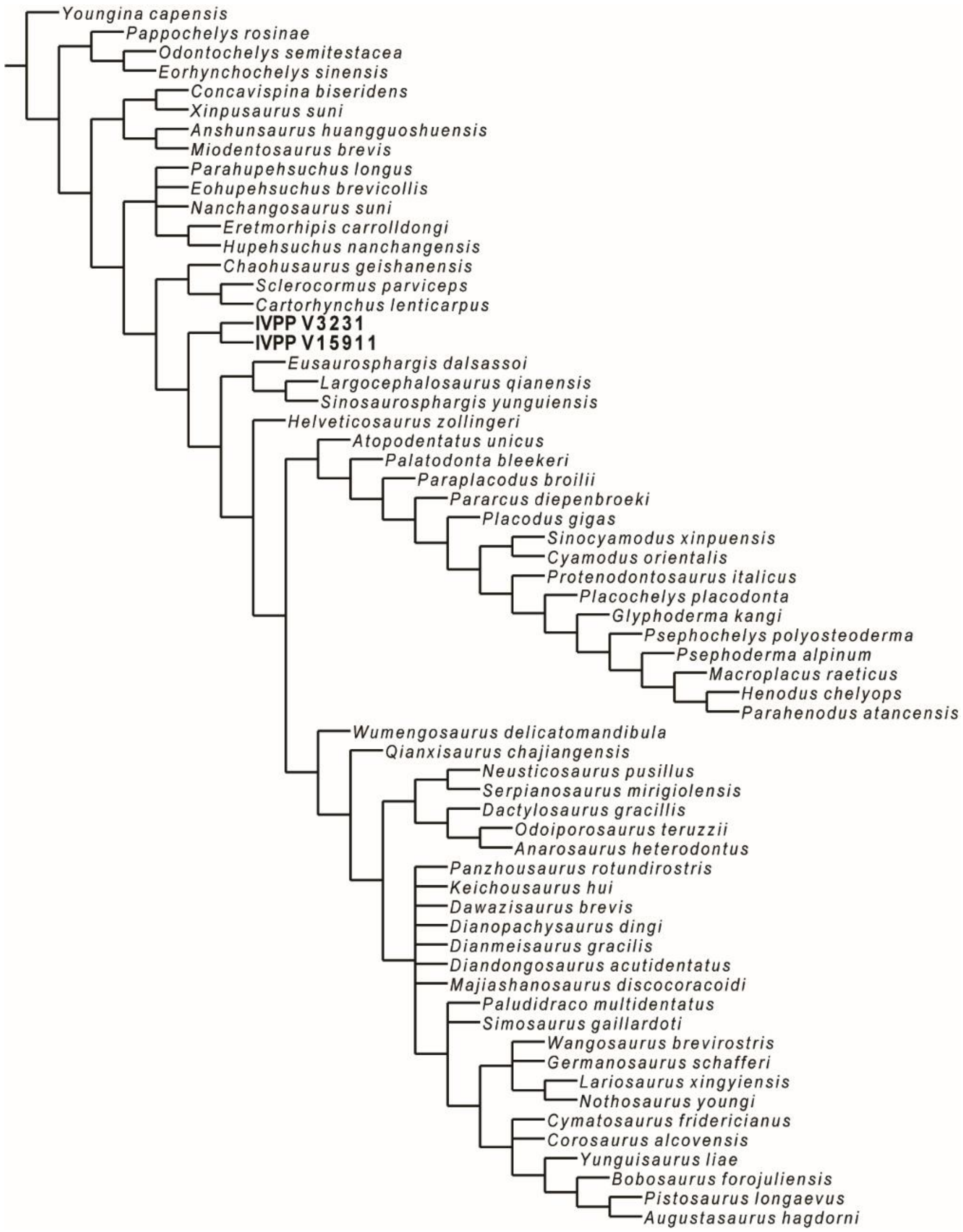


**Supplementary Fig. 5. Strict consensus tree based on our new data matrix recovering IVPP V 3231 and IVPP V 15911 as a single clade.**

62 taxa involved including *Youngina capensis* as the out group, basal pantestudines, hupehsuchians, ichthyosauriforms, thalattosaurians and all sauropterygiforms, 790 steps, 18 MPTs, CI: 0.301, RI: 0.700.

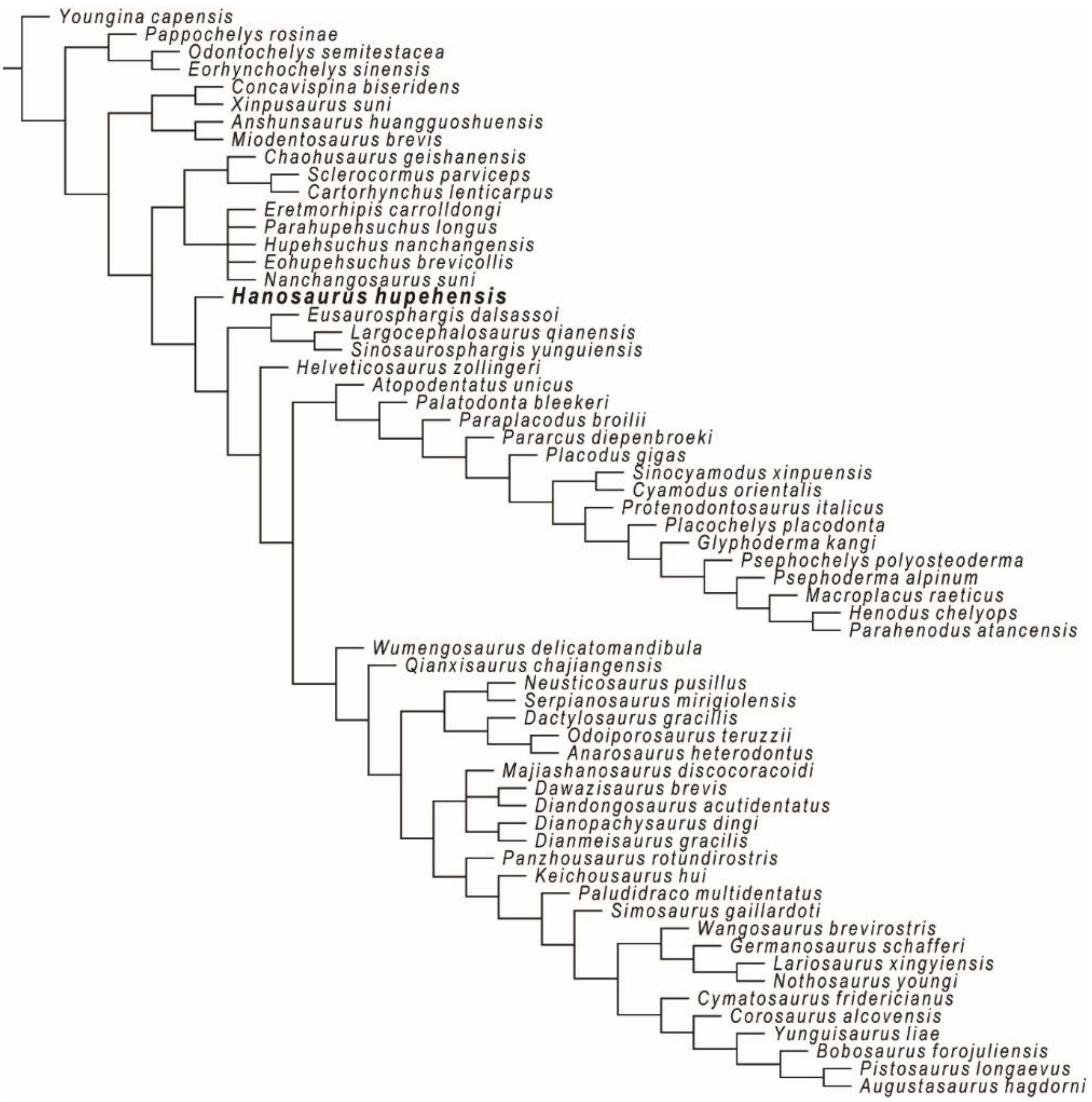


**Supplementary Fig. 6. Majority-rule consensus (50%) tree based on our new data matrix recovering *Hanosaurus hupehensis* as the basal-most taxon of sauropterygiforms.**

62 taxa involved including *Youngina capensis* as the out group, basal pantestudines, hupehsuchians, ichthyosauriforms, thalattosaurians and all sauropterygiforms, 790 steps, 100 MPTs, CI: 0.301, RI: 0.700.

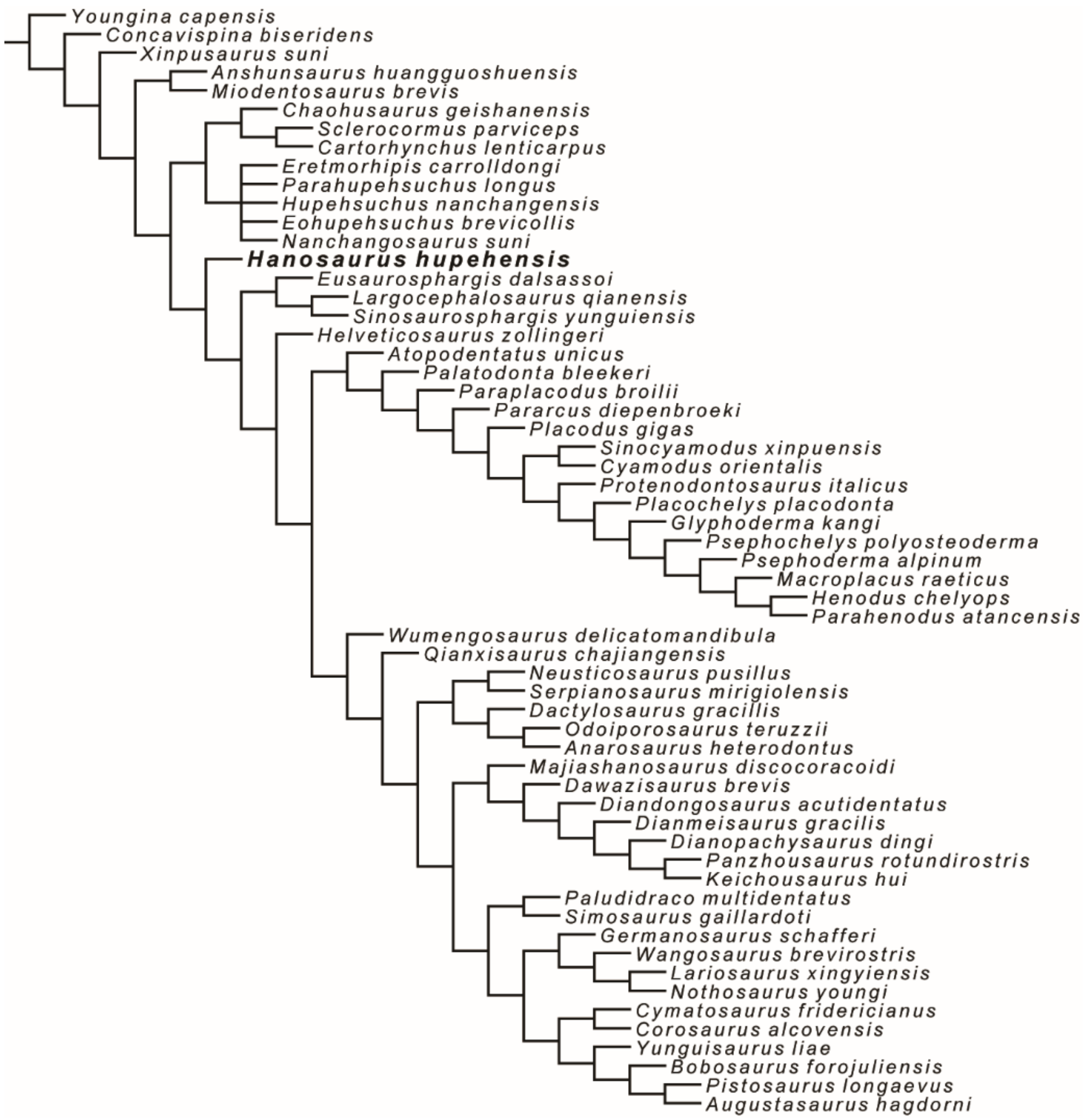


**Supplementary Fig. 7. Strict consensus tree based on part of our new data matrix recovering *Hanosaurus hupehensis* as the basal-most taxon of sauropterygiforms.**

59 taxa involved including *Youngina capensis* as the out group, hupehsuchians, ichthyosauriforms, thalattosaurians and all sauropterygiforms, but excluding basal pantestudines, 756 steps, 17 MPTs, CI: 0.315, RI: 0.718. Fig. 2 in the main text is mainly based on this result.

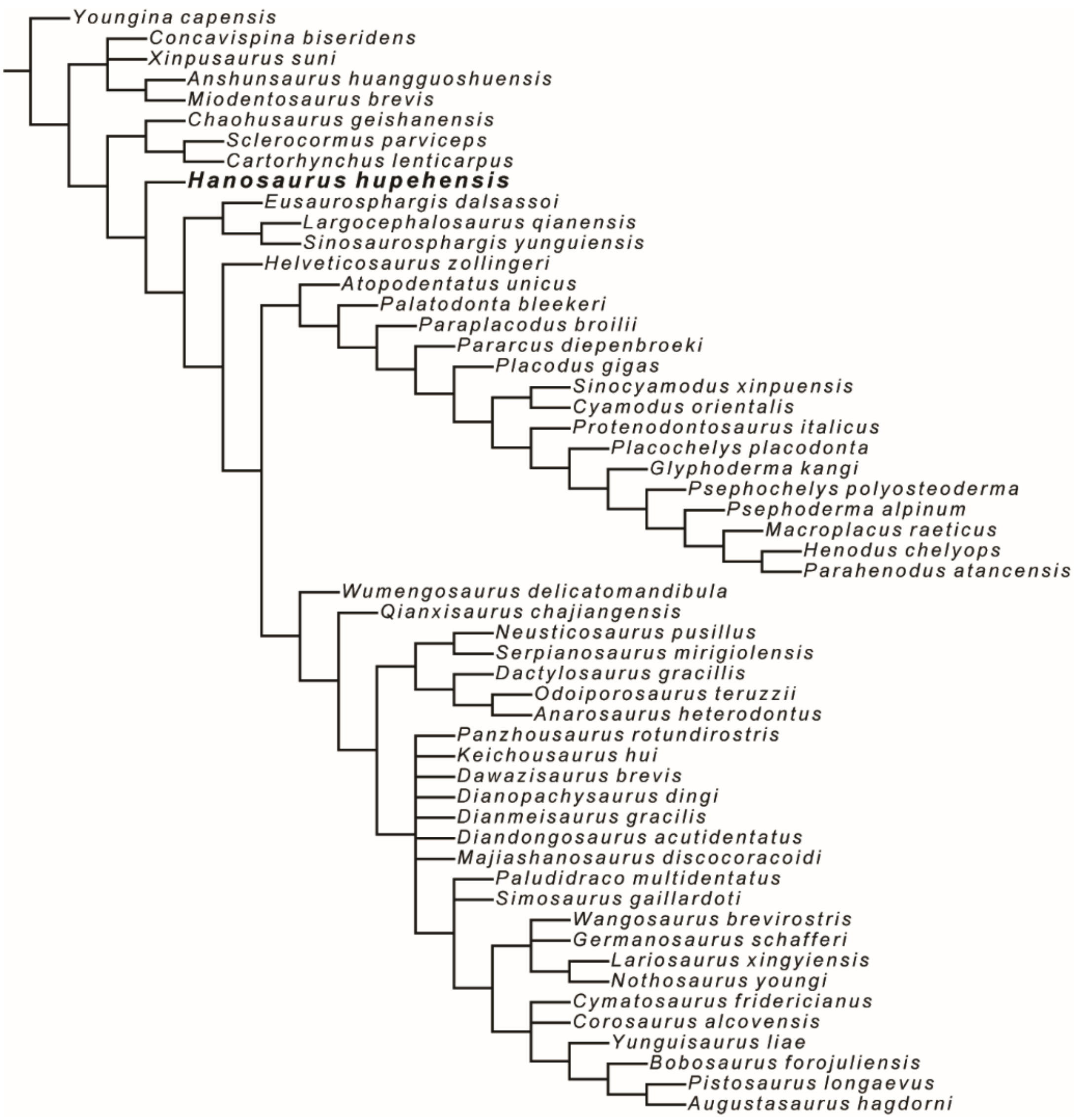


**Supplementary Fig. 8. Strict consensus tree based on part of our new data matrix recovering *Hanosaurus hupehensis* as the basal-most taxon of sauropterygiforms.**

54 taxa involved including *Youngina capensis* as the out group, ichthyosauriforms, thalattosaurians and all sauropterygiforms**,** but excluding basal pantestudines and hupehsuchians as unstable taxa in previous analyses, 724 steps, 12 MPTs, CI: 0.329, RI: 0.735.

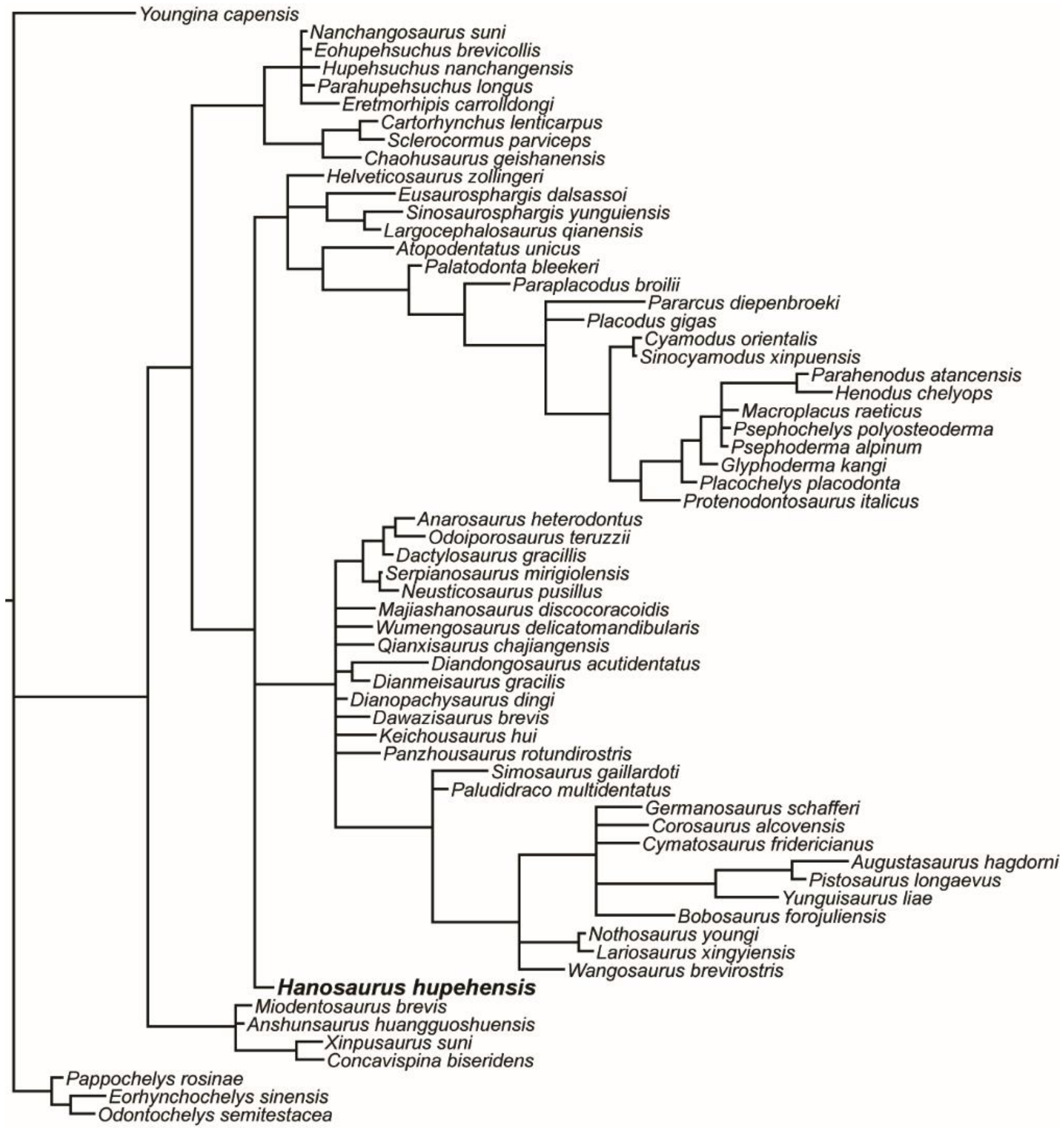


**Supplementary Fig. 9. Majority-rule consensus (50%) Bayesian tree based on our new data matrix recovering *Hanosaurus hupehensis* as the basal-most taxon of sauropterygiforms.**

62 taxa involved including *Youngina capensis* as the out group, basal pantestudines, hupehsuchians, ichthyosauriforms, thalattosaurians, and all sauropterygiforms.

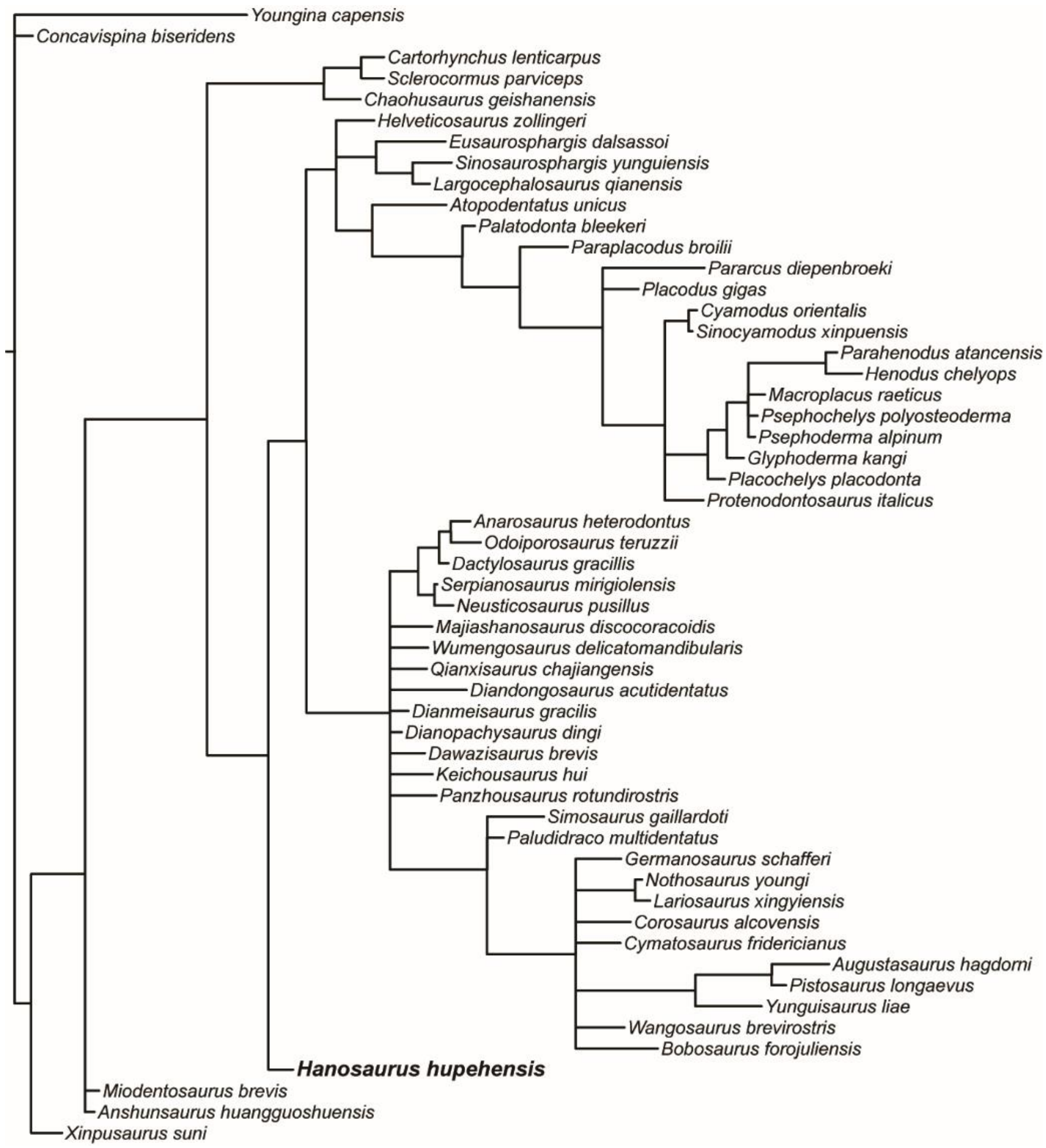


**Supplementary Fig. 10. Majority-rule consensus (50%) Bayesian tree based on part of our new data matrix recovering *Hanosaurus hupehensis* as the basal-most taxon of sauropterygiforms.**

54 taxa involved including *Youngina capensis* as the out group, ichthyosauriforms, thalattosaurians, and all sauropterygiforms**,** but excluding basal pantestudines and hupehsuchians as unstable taxa in previous analyses.

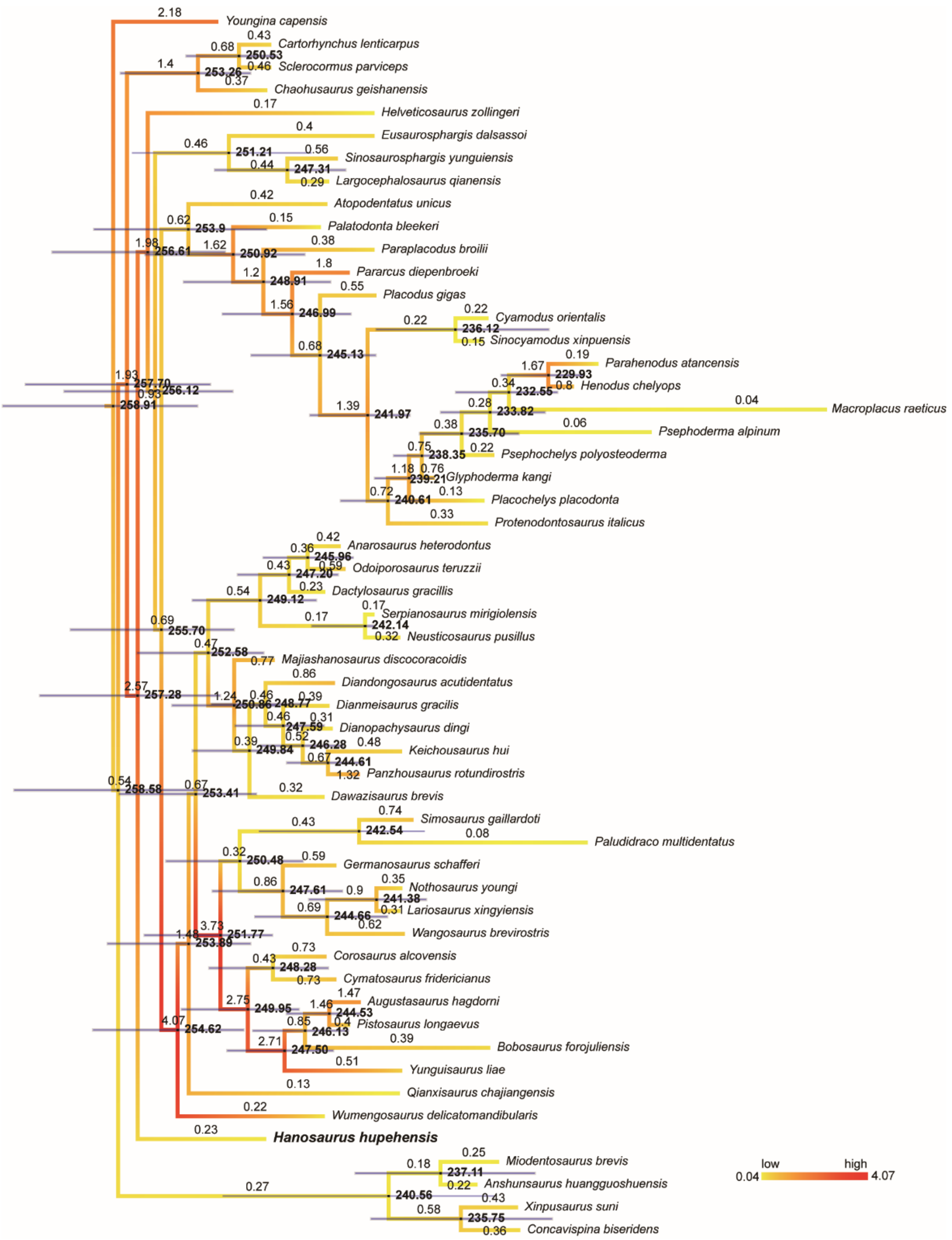


**Supplementary Fig. 11. Tip-dating result of Triassic marine reptiles.**

The topology of the cladogram is constrained based on our parsimony and bayesian results. Node labels are the medians of estimated ages, node bars reflect the 95% highest posterior density of ages; branch labels are the medians of evolutionary rates, branches are colored from yellow to red according to the evolutionary rate from low to high.

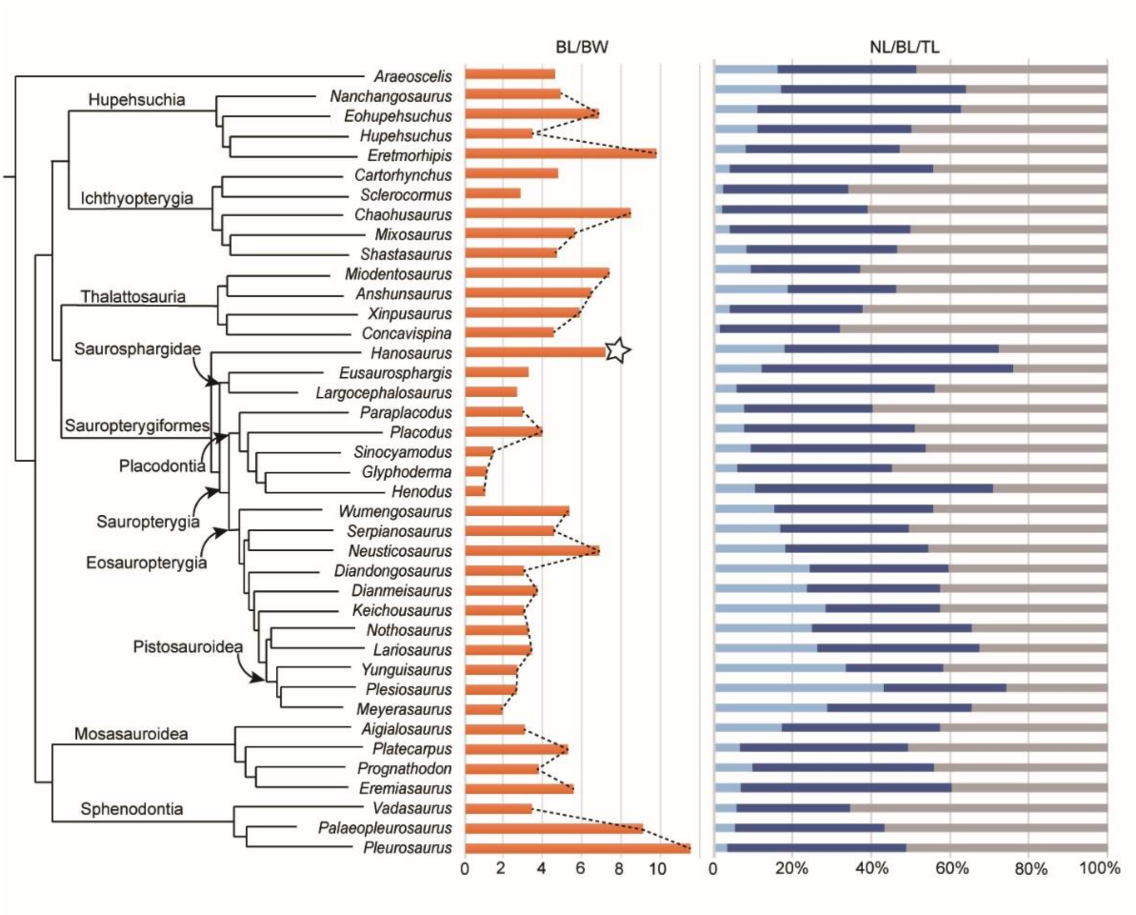


**Supplementary Fig. 12. Body shape and proportion comparisons Mesozoic marine reptiles.**

The phylogenetic topologies are illustrated based on our results and literature (Fig. 2, Supplementary Data 4). Orange bars indicate the ratios of body length to body width in each animal with *Hanosaurus* marked, which generally increase from the terrestrial (paraxial locomotion) to coastal (axial locomotion) marine taxa and decrease from coastal (axial locomotion) to pelagic (paraxial locomotion) marine taxa in each group. Light blue, dark blue, and gray bars reflect the proportions of neck length, body length, and tail length respectively among the sums of these three regions (see abbreviations in Materials and Methods).

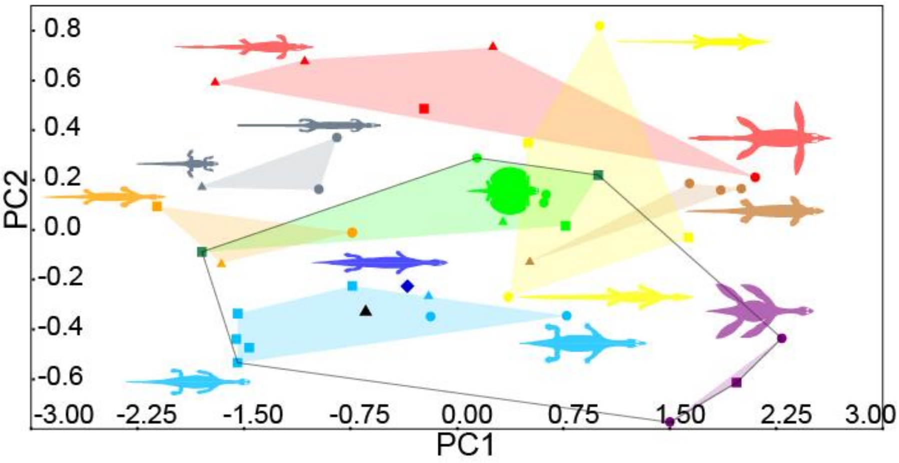


**Supplementary Fig. 13. Morphospace based on principal components PC1 and PC2.**

Black lines outline the area occupied by sauropterygiform reptiles (see the detailed explanation of colors and dot shapes in Fig. 3).

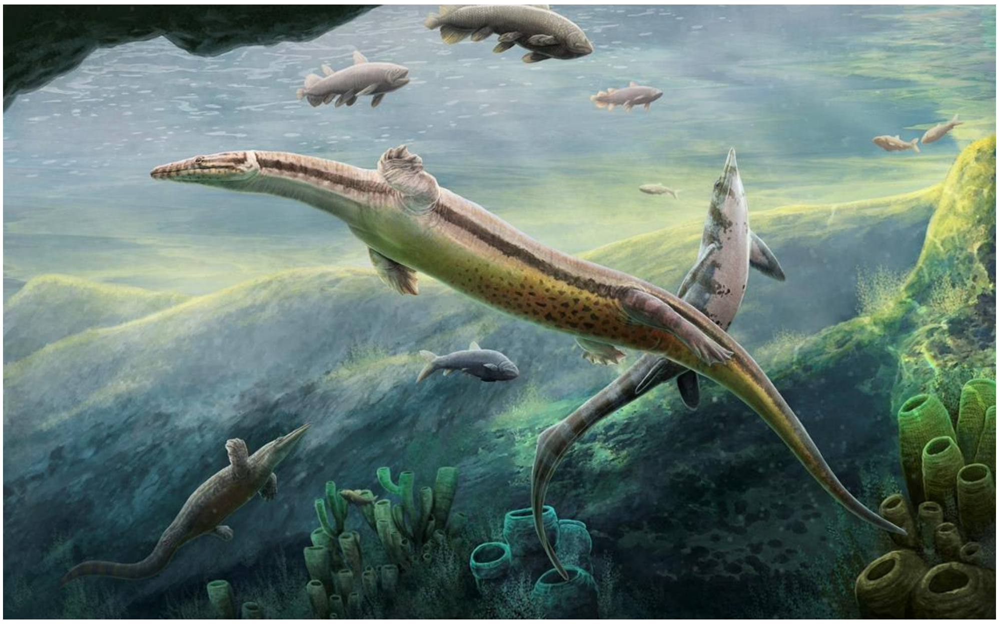

**Supplementary Fig. 14. Marine reptiles of the shallow marine habitat in the Early Triassic, Hubei, China.**

Marine reptiles include *Hanosaurus hupehensis* (middle), *Chaohusaurus zhangjiawanensis* (right upper), *Nanchangosaurus suni* (left lower) surrounded by fishes and sponges. The long-trunk and short-limbs were convergently obtained in these reptiles showing axial undulatory locomotion. Reconstructed by Gabriel Ugueto.
